# DRP1-mediated regulation of mitochondrial dynamics determines the apoptotic response upon embryonic differentiation

**DOI:** 10.1101/835751

**Authors:** Barbara Pernaute, Juan Miguel Sánchez Nieto, Salvador Pérez-Montero, Aida di Gregorio, Ana Lima, Katerina Lawlor, Sarah Bowling, Gianmaria Liccardi, Alejandra Tomás, Pascal Meier, Guy A. Rutter, Ivana Barbaric, Tristan A. Rodríguez

## Abstract

The changes that drive differentiation create a large potential for the emergence of abnormal cells that need to be removed before they contribute to further development or the germline. This removal is in part achieved by cells becoming hypersensitive to death upon exit of naïve pluripotency. What causes this change in apoptotic response is unknown. Here we identify that it is controlled by the regulator of mitochondrial dynamics DRP1. We show that in mouse, naïve pluripotent cells have fragmented mitochondria due to high DRP1-mediated fission, but upon differentiation, DRP1 activity decreases, inducing mitochondria to fuse and form complex networks. We demonstrate that this decrease in DRP1 activity lowers the apoptotic threshold, as mutation of DRP1 increases the sensitivity to cell death and its over-expression protects against apoptosis. Together, our findings highlight how regulation of mitochondrial dynamics allows cells to adapt their apoptotic response to the changing environment of differentiation.

## Introduction

The appropriate regulation of apoptosis during embryonic development is essential for maintaining the right balance between the elimination of suboptimal cells and the availability of sufficient cell numbers to sustain embryo growth^1^. The maintenance of this balance is most acutely evident in the lead up to gastrulation, when the mouse epiblast not only undergoes the process of germ layer specification, but also displays a significant expansion in cell numbers, from 150 at E5.5 to 15,000 cells at E7.5^2^. This rapid proliferation of cells is concomitant with the substantial cellular changes that accompany gastrulation, most notably extinction of the pluripotency network and activation of differentiation genes in a lineage-specific manner. Any abnormal or mis-specified cell, which cannot perform these changes appropriately, needs to be removed from the embryo to prevent them from contributing to further development or the germline, which is also specified around E6.5^3^. Hence, a wave of cell death takes place in the mouse embryo at E6.5, most likely reflecting the elimination of these abnormal cells^4^, ^5^.

One way by which the embryo facilitates the elimination of aberrant cells is through lowering the apoptotic threshold as it proceeds from a pre-implantation to an early post-implantation stage of development. Indeed, whilst low doses of UV irradiation do not induce apoptosis in mouse pre-implantation embryos, these same low doses lead to a strong apoptotic response in the post-implantation epiblast^6^. Similarly, although cells with a range of genetic defects, including chromosome fragmentation or chromosome mis-segregation survive pre-implantation development, they are efficiently eliminated by apoptosis during early post-implantation development^7–10^. Subsequently, it has been proposed that the post-implantation epiblast is primed for death, and therefore hyper-responsive to apoptotic signals^6^, ^11^.

The mitochondrial apoptotic pathway is tightly regulated by a balance between pro- and anti-apoptotic factors belonging to the BCL-2 family. The binding of the anti-apoptotic proteins (e.g BCL-2, BCL-XL, A1 or MCL-1) to their pro-apoptotic BCL-2 family counterparts (e.g. BIM, BID, PUMA, NOXA or BAD) prevents the induction of apoptosis. When this balance is lost, BIM or BID bind to the apoptotic effector molecules BAX and BAK, which induce mitochondrial outer membrane permeabilization (MOMP) and caspase activation^12^. We have previously shown that, in the post-implantation epiblast, this balance is at least in part maintained by microRNAs (miRNAs) of the miR-20, miR-92, and miR-302 families. These miRNAs target *Bim* (*Bcl2l11*) and maintain it in a state that is poised for activation^11^. However, it is worth noting that mutation of *Bim* does not prevent the endogenous wave of cell death occurring during early mouse development, suggesting that other factors must be contributing to making the epiblast hypersensitive to death signals^11^.

The balance between mitochondrial fusion and fission, termed mitochondrial dynamics, has emerged over the last few years as an important regulator of the apoptotic response. Mitochondrial dynamics are facilitated by proteins such as MFN1, MFN2 and OPA1, that promote fusion and DRP1 or MFF, that promote fission^13–15^. During cell death, remodelling of the cristae network regulated by OPA1^16^ and DRP1^17^, ^18^ is required for release of cytochrome C and subsequent apoptosis. DRP1-induced mitochondrial fragmentation is also coupled to the later stages of apoptosis^19–22^. Finally, MFN1-induced fusion has been shown to promote a mitochondrial size that is permissive for BAX function^23^. Therefore, the regulators of mitochondrial dynamics play a variety of different roles in the apoptotic process. In the developing embryo, mitochondrial shape undergoes substantial changes around the time of implantation^24^. During pre-implantation stages mitochondria are round and have a low cristae density^25^, but later in development mitochondria elongate and cristae density increases^24^. These observations raise the possibility that changes in mitochondrial dynamics may play roles in regulating the changes in apoptotic threshold that occur during early development.

Here we have identified that a change in mitochondrial dynamics increases the sensitivity to cell death signals in cells of the post-implantation epiblast. For this, we combined experiments in the embryo, with studies using embryonic stem cells (ESCs), which capture the naïve pluripotent state found in the pre-implantation epiblast, and in epiblast stem cells (EpiSCs), which resemble the primed pluripotent post-implantation epiblast^26^. We find that although primed pluripotent cells show high mitochondrial apoptotic priming, their increased sensitivity to cell death is not due to differential expression of members of the BCL-2 pro- and anti-apoptotic family. In contrast, we observed that decreased mitochondrial fission correlates with the readiness of cells to undergo apoptosis. Furthermore, we demonstrate that manipulating DRP1 activity is sufficient to change the apoptotic threshold of pluripotent cells. Together, these results demonstrate that changes in mitochondrial dynamics influence the apoptotic priming status of cells and contribute to the elimination of aberrant cells during early embryonic development.

## Results

### Pluripotent cells become hypersensitive to cell death upon exit of naïve pluripotency

We set out firstly to characterise the apoptotic sensitivity of the different pluripotent cell types found during early mouse development. Previous studies have shown that whilst cells from the pre-implantation embryo are relatively resistant to low doses of UV radiation, the early post-implantation cells respond to the same low doses of UV by undergoing apoptosis^6^. In line with these findings, we observed no overt apoptosis in E3.5 mouse embryos treated with the DNA damage-inducing drug etoposide for 1.5h, but we detected a strong apoptotic response when the etoposide treatment was applied to E6.5 embryos (Supplementary Figure 1A-B).

**Figure 1.**
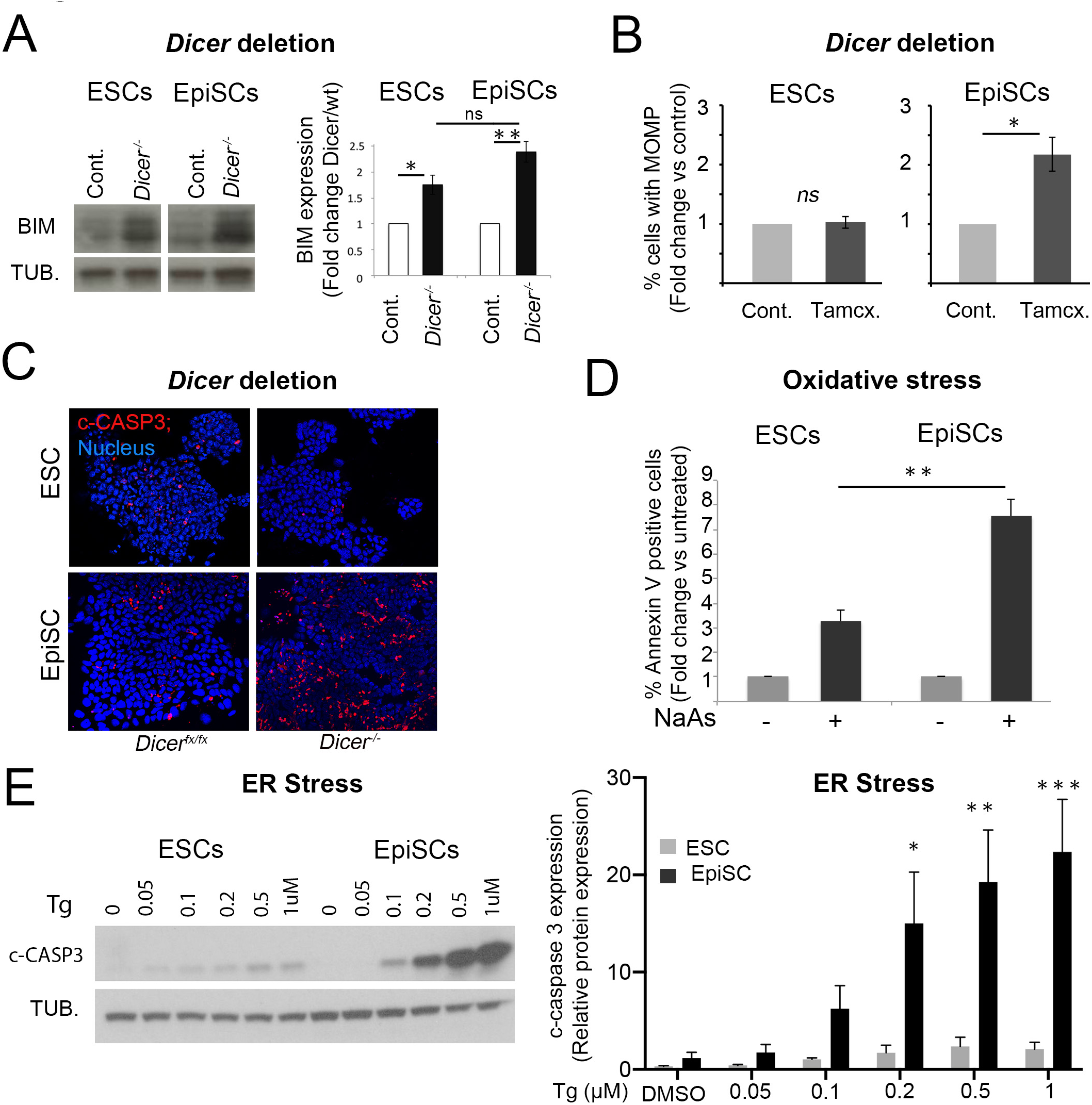
The apoptotic threshold decreases upon pluripotent stem cell differentiation. **A.** Increase in BIM expression upon *Dicer* deletion in EpiSCs (day 6 post-deletion) and ESCs (day 7 post-deletion). Representative western blot and western blot quantification relative to α-TUBULIN (TUB.). **B.** Change in % cells with mitochondrial outer membrane permeabilization (MOMP) upon *Dicer* deletion in EpiSCs (day 5 post-deletion) and ESCs (day 6 post-deletion) relative to un-deleted cells as measured by DiOC6 staining. **C.** Cleaved-CASPASE 3 (c-CASP3) immunostaining in ESCs and EpiSCs upon *Dicer* deletion at day 5 and 6 respectively. **D.** Change in % Annexin V positive cells after 16h treatment with 1µM sodium arsenite relative to untreated cells. **E**. Cleaved-CASPASE 3 levels in ESCs and EpiSCs upon treatment for 16h with increasing concentrations of thapsigargin. Representative western blot and graph showing average western blot quantification relative to α-TUBULIN. Average of 3 (A, E) or 4 (B, D) experiments +/-SEM is shown. Students T-Test (A-B and D) or 2-way ANOVA with Šidák correction (E) *p<0.05, **p<0.01, ***p<0.001. ns= not significant.

We next tested whether the observed differential sensitivity to apoptosis is also apparent in ESCs and EpiSCs, the two *in vitro* counterparts of the pre- and post-implantation epiblast^26^. We have previously shown that loss of miRNAs leads to an upregulation of BIM expression and a consequent induction of apoptosis in EpiSCs^11^. To test if miRNAs also regulate apoptosis in ESCs, we induced *Dicer* deletion in *Dicer*^*Fx/Fx*^ ESCs by tamoxifen administration, as we had previously done for EpiSCs^11^. This led to miRNA depletion (Supplementary Figure 1C) and an up-regulation of BIM expression similar to that found in EpiSCs (Figure 1A). However, in contrast to what was seen in EpiSCs, the increase in BIM expression did not cause apoptosis in ESCs (Figure 1B-C and Supplementary Figure 1D). This suggests that mouse ESCs are more resistant to apoptosis than EpiSCs.

Importantly, we found that the differential apoptotic response of ESCs and EpiSCs was also evident upon exposing these cells to different sources of stress. Induction of oxidative stress with 1µM sodium arsenite for 16 hours produced a 3-fold increase in Annexin V positive ESCs, but a 7-fold increase in Annexin V positive EpiSCs (Figure 1D). Similarly, induction of endoplasmic reticulum (ER) stress with increasing doses of thapsigargin for 16 hours led to a small change in the basal levels of cleaved Caspase 3 in ESCs, whereas the same treatment induced a robust apoptotic response in EpiSCs (Figure 1E). Overall, these results indicate that EpiSCs exhibit an increased sensitivity to apoptosis when compared to ESCs, thus recapitulating the features observed in their *in vivo* counterparts.

### The mitochondrial apoptotic pathway is primed for cell death in primed pluripotent stem cells

We next addressed if the differences in apoptotic response between naïve and primed pluripotent cells were reflected at the level of the mitochondrial apoptotic pathway. This pathway is regulated by the relative expression of pro-apoptotic and anti-apoptotic BCL-2 family members^12^. This balance can be artificially changed, for example by inhibiting BCL-2/XL activity with BH3 mimetics such as ABT-737. We observed that 24h treatment of EpiSCs with ABT-737 led to a strong increase in the percentage of cells displaying (1) MOMP (Supplementary Figure 1E), (2) Annexin V positivity (Figure 2A) and (3) cleaved caspase-3 expression (Figure 2B). In contrast, treatment with ABT-737 failed to induce apoptosis in ESCs, even at the highest concentration used (Figure 2A-B and Supplementary Figure 1E-F). These findings are strengthened by the observation that ABT-737 leads to cytochrome C release from the mitochondria in EpiSCs, but not ESCs (Figure 2B). To test the physiological significance of these observations, we treated pre- and post-implantation mouse embryos with 2μM ABT-737 for 1.5h. Similar to what we observed *in vitro*, we found that this treatment induced apoptosis in epiblast cells of E6.5 embryos, but not in inner cell mass cells of E3.5 embryos (Figure 2C and Supplementary Figure 1F).

**Figure 2.**
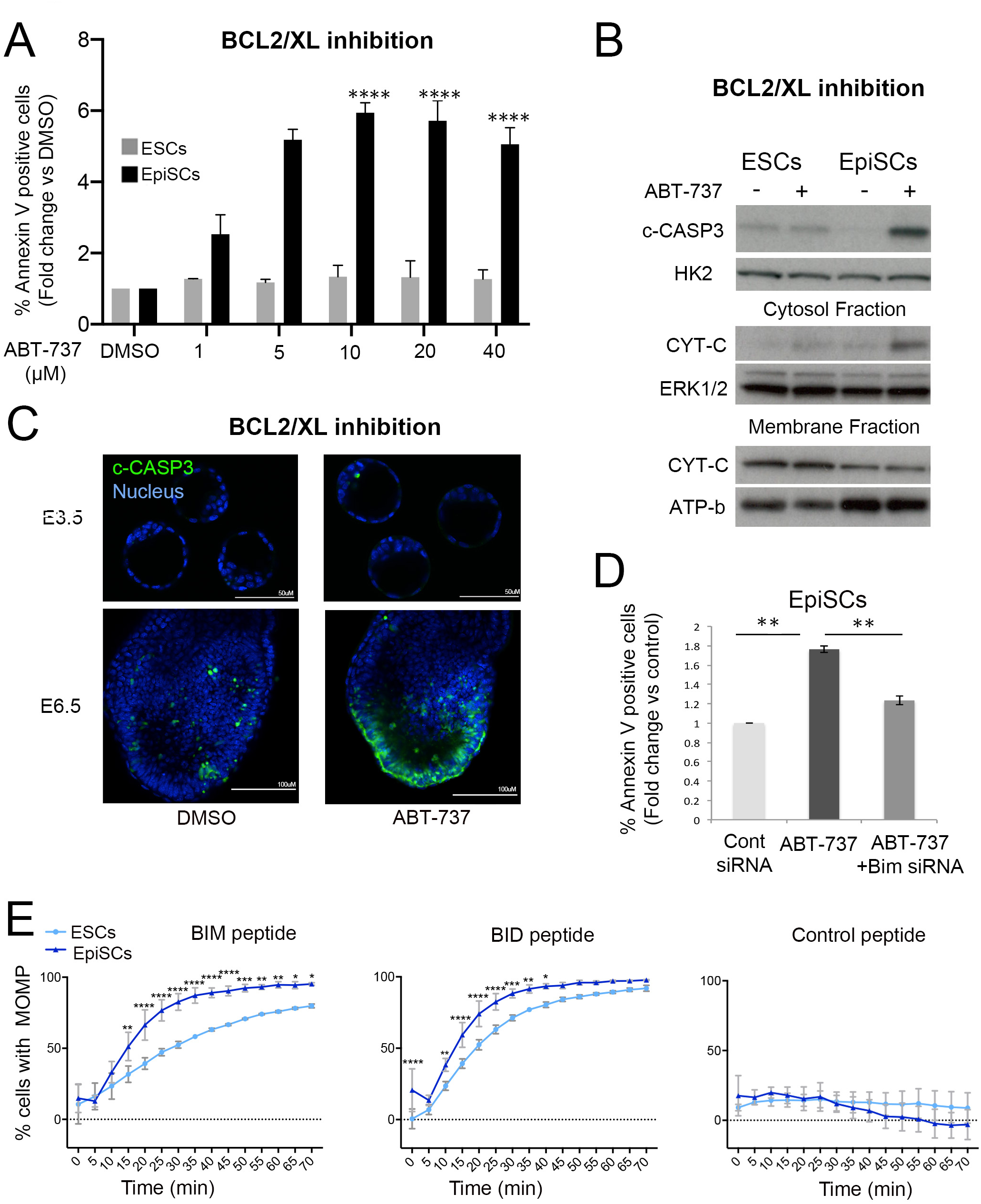
Enhanced activation of the mitochondrial apoptotic pathway in primed pluripotent cells. Apoptosis quantified as % Annexin V positive ESCs and EpiSCs treated for 24h with increasing concentrations of ABT-737. Fold change versus dmso treated cells is shown. **B**. Cleaved-CASPASE 3 (c-CASP3) levels relative to HEXOKINASE 2 (HK2) in ESCs and EpiSCs treated for 24h with 5µM ABT-737. CYTOCHROME C (CYT-C) levels in the cytosolic and membrane fractions in ESCs and EpiSCs treated with 5µM ABT-737 for 24h relative to ERK1/2 and ATP-b. **C**. Cleaved-CASPASE 3 immunostaining in E3.5 and E6.5 embryos treated with dmso (E3.5 n= 3; E6.5 n=5) or 2µM ABT-737 (E3.5 n= 3; E6.5 n=5) for 1.5h. **D**. Fold change in % Annexin V positive EpiSCs transfected with control siRNA or Bim siRNA and treated with dmso or 5µM ABT-737 for 24h. **E**. % membrane depolarization in ESCs and EpiSCs treated with with either the BIM (10µM), BID (10µM) or control (10µM) peptides for the indicated amounts of time. Average of 3 (A, E) or 4 (B) experiments +/-SEM (D) or +/-SD (E) is shown. (D) Student T-Test or (A) 2-way ANOVA with Šidák correction *p<0.05, **p<0.01 or ****<0.00001.

Next, we determined whether the enhanced sensitivity of EpiSCs to BCL2/BCLXL inhibition was dependent on BIM. For this we transfected previously tested siRNAs targeting Bim^11^ into EpiSCs and treated them with ABT-737. We observed that this was sufficient to suppress the apoptotic response induced by BCL2/BCL-XL inhibition (Figure 2D), further highlighting the importance of BIM for the apoptotic response of EpiSCs. Given that the differences in apoptotic response between ESCs and EpiSCs appear to be mediated by the mitochondrial apoptotic pathway, we measured the level of mitochondrial apoptotic priming of these cells by analysing the kinetics of mitochondrial membrane depolarization induced by BIM and BID BH3 peptides^27^. We observed that the depolarization of the mitochondrial membrane potential was significantly more pronounced in EpiSCs than in ESCs for both these peptides (Figure 2E). Together these results indicate that primed pluripotent cells have a lower apoptotic threshold than naïve cells due to enhanced sensitivity of the mitochondrial pathway.

### The relative expression of pro- and anti-apoptotic BCL2 family members does not explain the different apoptotic threshold of ESCs and EpiSCs

To address if differences in the relative expression of pro- and anti-apoptotic BCL2 family members underpin the different apoptotic response of ESCs and EpiSCs, we compared the expression of these proteins in each cell type. We observed no significant difference in the expression of the pro-apoptotic activator proteins PUMA, BIM, BID or BAD in either whole cell or mitochondrial extracts between these two cell types (Figure 3A). Similarly, we did not observe any difference in the expression of the pro-apoptotic effector proteins BAX or BAK (Figure 3B). In contrast, when anti-apoptotic proteins were analysed, we found little difference in the expression of BCL-XL, MCL1 or A1, but significantly higher expression of BCL-2 in EpiSCs, in both whole cell and mitochondrial extracts (Figure 3C). Whilst a higher expression of anti-apoptotic protein expression may seem counterintuitive given the enhanced sensitivity to death of EpiSCs, this elevated BCL-2 expression is likely part of the adaptation of these cells to their low apoptotic threshold^28^.

**Figure 3.**
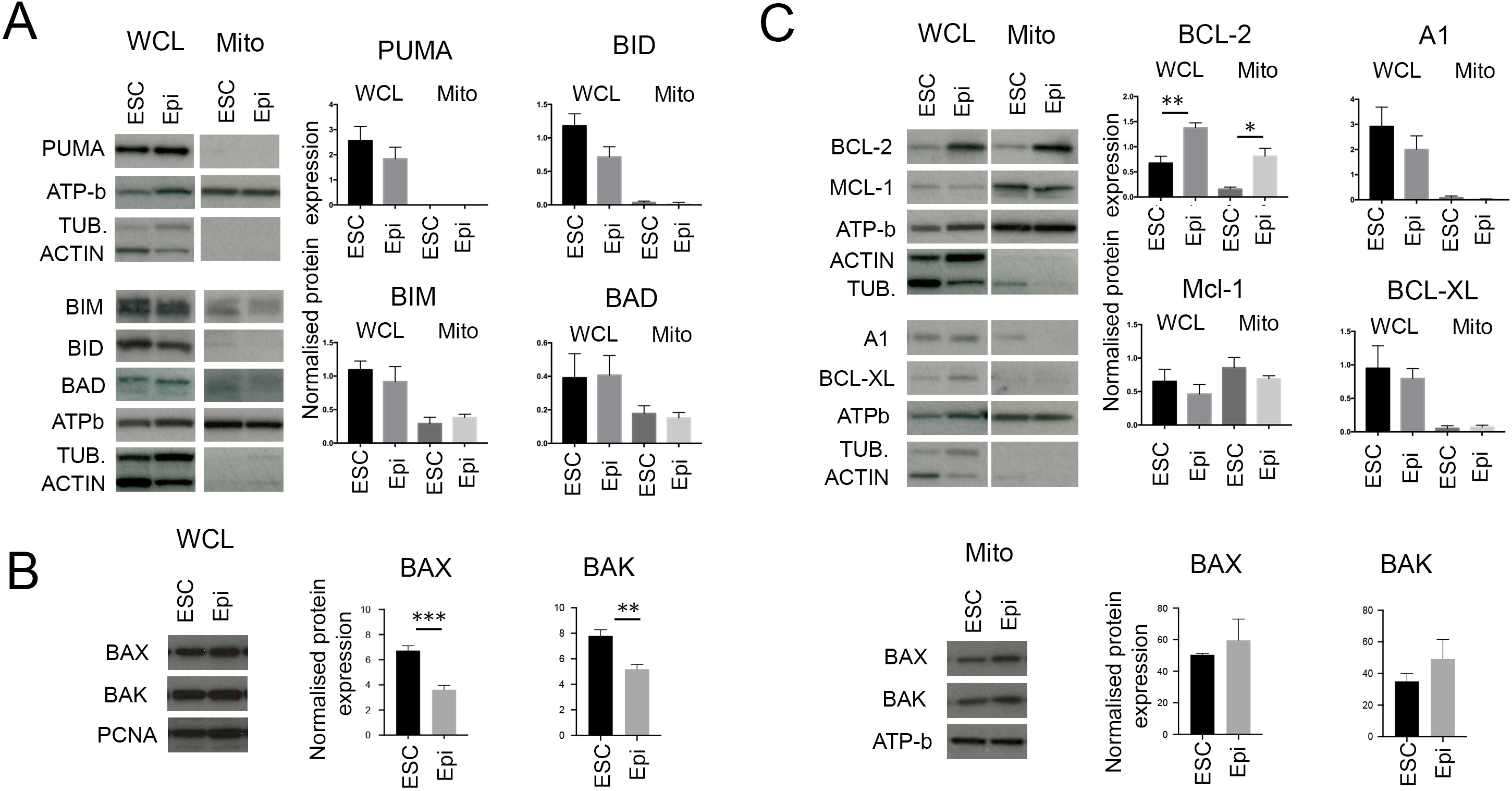
Expression levels of BCL-2 apoptotic factors in pluripotent cells. **A.** Levels of pro-apoptotic factors in whole cell lysate and mitochondrial extract of ESCs an EpiSCs (Epi). Graphs show protein level normalized against α-TUBULIN (TUB.) or β-ACTIN in whole cell lysate and ATP-b in mitochondrial extract. **B.** Levels of apoptosis effectors in whole cell lysate and mitochondrial extract of ESCs an EpiSCs. Graphs show protein level normalized against αTUBULIN, βACTIN or PCNA for the whole cell lysates and ATP-b for the mitochondrial extracts. **C**. Levels of anti-apoptotic factors in whole cell lysate and mitochondrial extract of ESCs an EpiSCs. Graphs show protein level normalized against αTUBULIN or βACTIN in whole cell lysate and ATP-b in mitochondrial extract. Average of 4 independent experiments +/-SEM is shown. Students T-Test *p<0.05, **p<0.01, ***p<0.001

To determine if the balance of expression of pro- and anti-apoptotic factors changes differently in ESCs and EpiSCs upon induction of apoptosis, we analysed the expression of key BCL-2 family members after treating these cells with 5µM ABT-737 for 24 hours. In these experiments we analysed the expression of the anti-apoptotic BCL-2 family members as well as BIM expression, as BIM is required for the apoptotic response to ABT-737 (Figure 2D). However, we observed no significant change in the levels of expression of BCL-2, BCL-XL, A1, MCL-1 or BIM between ABT-737 treated samples and controls in either whole cell or mitochondrial extracts of ESCs or EpiSCs (Supplementary Figure 2A-B). This suggests that the induction of apoptosis does not significantly shift the balance of pro- and anti-apoptotic BCL-2 family expression. Together, these results indicate that the relative expression of BCL-2 family proteins is not the cause of the higher sensitivity to apoptosis of primed pluripotent cells.

### High levels of mitochondrial fission are observed in naïve pluripotent cells when compared to primed cells

Mitochondrial dynamics have been suggested as a mechanism that contributes to the regulation of the mitochondrial apoptotic threshold^13^. The mitochondria of ESCs and pre-implantation embryos have been shown to be rounded^25^ and to elongate upon differentiation^29^, ^30^. Analysis of mitochondrial morphology in ESCs revealed rounded doughnut shaped mitochondria in agreement with previous studies^30^. In contrast, the mitochondria of EpiSCs were more elongated (Figure 4A). Importantly, these differences were also seen *in vivo*, with E3.5 inner cell mass cells and E4.5 epiblast cells having rounded mitochondria and the mitochondria of E6.5 epiblast cells being elongated and forming networks (Figure 4B).

**Figure 4.**
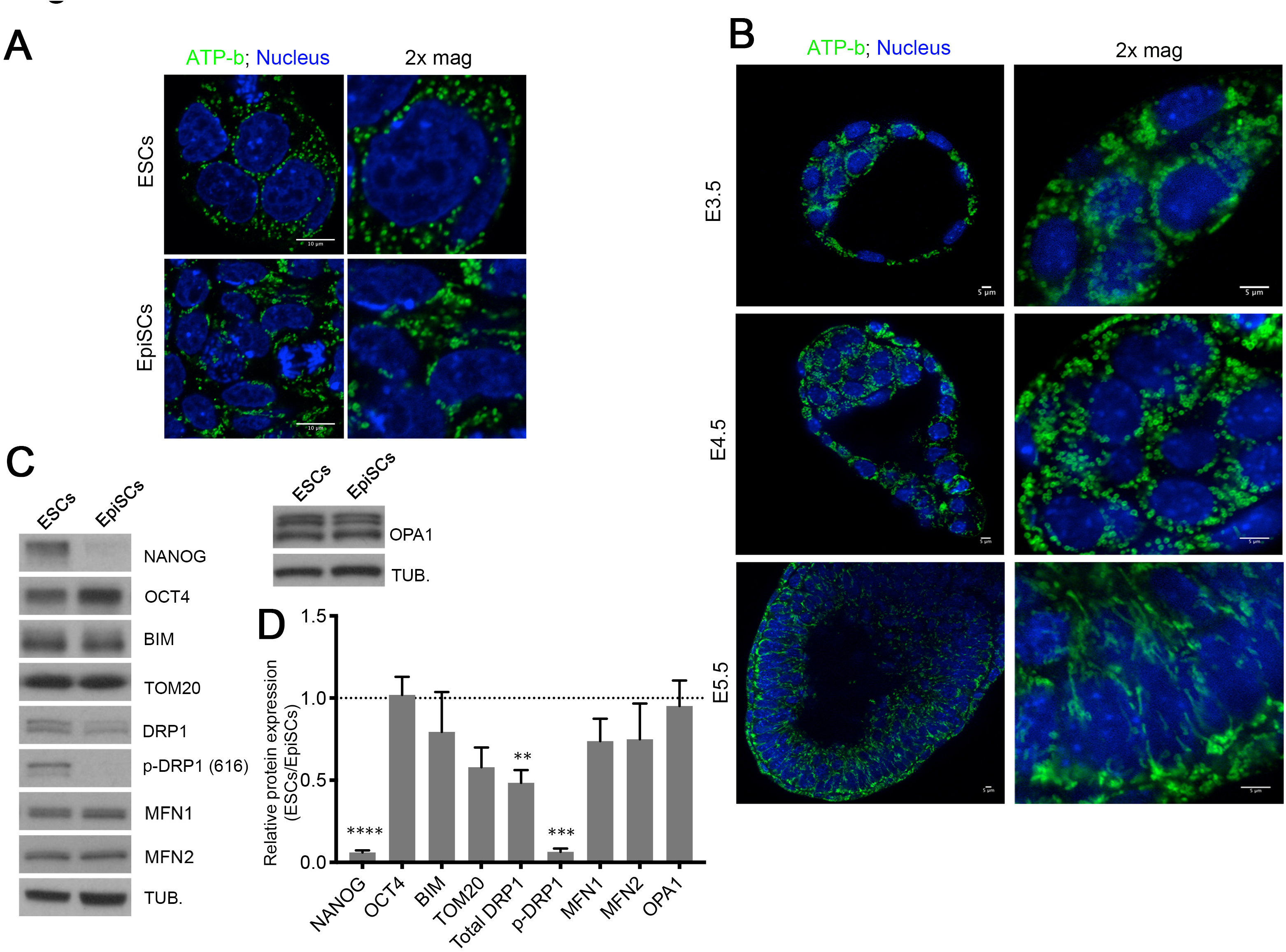
Mitochondria fuse to form complex networks upon differentiation. **A.** ATP-b immunostaining showing mitochondrial morphology in ESCs and EpiSCs. **B**. ATP-b immunostaining showing mitochondrial morphology in E3.5, E4.5 and E6.5 mouse embryos. 2x magnification over ICM/Epiblast area. **C**. Basal levels of mitochondrial fusion and fission proteins in ESC and EpiSCs. **D.** Mitochondrial fusion and fission protein levels in EpiSCs versus ESCs. Protein levels are normalized against α-TUBULIN (TUB.). Graph shows the average from 3 independent experiments +/-SEM is shown. 2-way ANOVA with Šidák correction **p<0.01, ***p<0.001 or ****<0.00001.

The differences in mitochondrial shape between naïve and primed pluripotent cells suggest that these cell types have different mitochondrial dynamics. We therefore studied the expression of fusion and fission regulators in ESCs and EpiSCs. While we observed no difference in the expression of the fusion regulators MFN1, MFN2 or OPA1 between these cell types, we noticed that ESCs had significantly higher levels of total DRP1 and p-DRP1 (S616), a phosphorylation event that induces fission^31^ (Figure 4C-D). This indicates that naïve pluripotent cells have higher fission activity. To determine if the higher p-DRP1 expression is responsible for the rounded mitochondrial shape of ESCs, we knocked-out *Drp1* by CRISPR-Cas9. Although *Drp1*^*−/−*^ ESCs remained pluripotent, their mitochondria became elongated and hyperfused (Figure 5A-B and Supplementary Figure 3A). These results indicate that DRP1-induced mitochondrial fission is required for the fragmented mitochondrial shape observed in ESCs.

**Figure 5.**
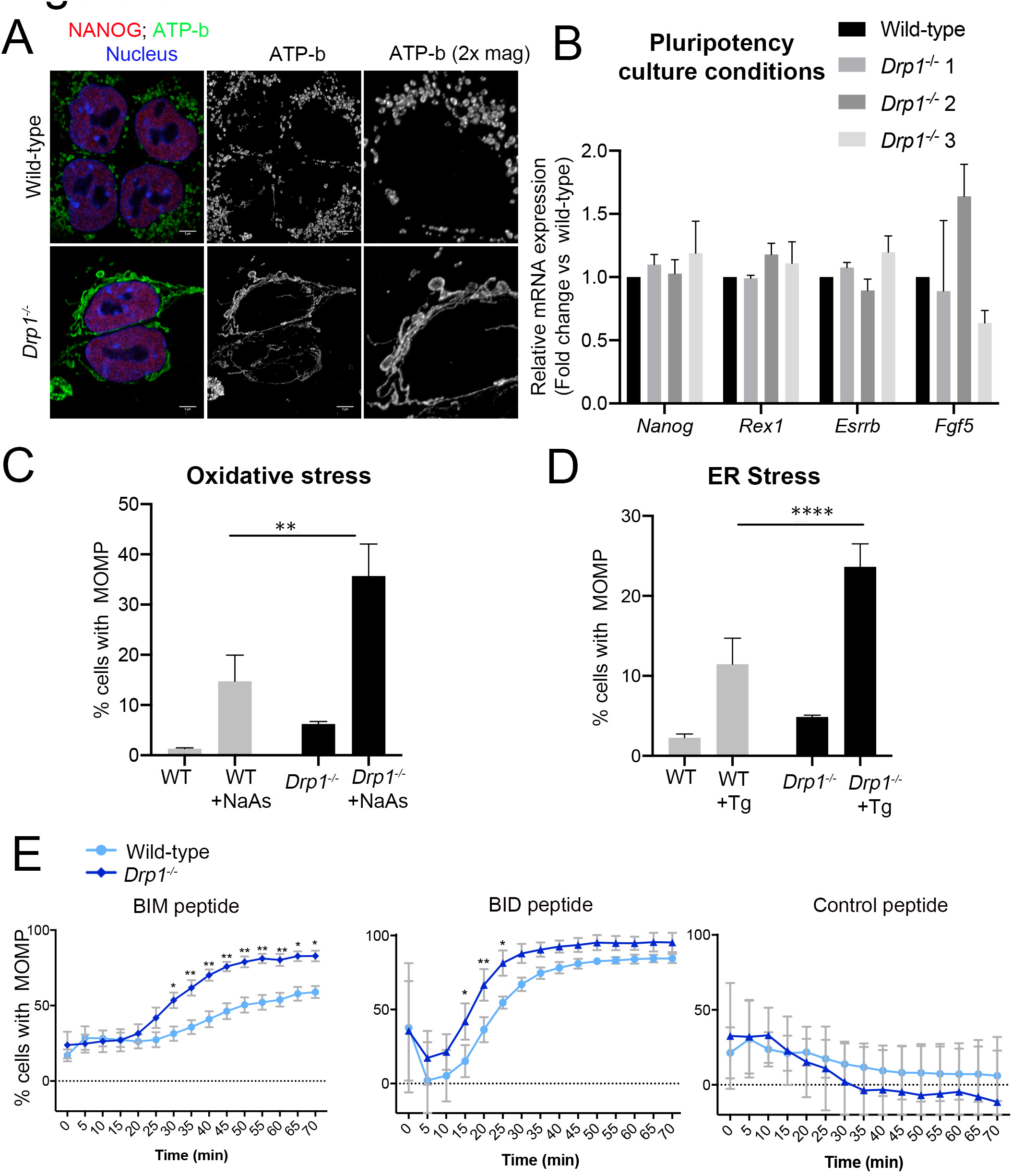
*Drp1* deletion facilitates early apoptotic events. **A.** ATP-b and NANOG immunostaining showing mitochondrial morphology in wild type and *Drp1*^*−/−*^ ESCs. **B**. Quantitative RT-PCR showing gene expression levels of naïve and primed pluripotency markers in wild-type and *Drp1*^*−/−*^ ESCs. Gene expression normalized against *Gapdh*. **C**. % of cells with MOMP detected by TMRM staining in wild-type and *Drp1*^*−/−*^ ESCs untreated or treated with 1µM sodium arsenite for 16h or **D** 1µM Thapsigargin for 16h. Data normalized against wild-type cells. **E**. % cells with MOMP in ESCs and EpiSCs treated with with either the BMI (0.5µM), BID (2.5µM) or control (1µM) peptides for the indicated amounts of time. Average of 3 (B, C), 4 (D) or 7 (E) independent experiments +/-SEM (C, D) or +/-SD (E) is shown. 2-way ANOVA with Šidák correction *p<0.05, **p<0.01 or ****<0.00001.

### DRP1 plays a key role in determining the apoptotic threshold of naïve and primed pluripotent cells

To evaluate whether DRP1 levels determine the apoptotic threshold of pluripotent cells we first analysed the effects of loss of *Drp1*. In the first instance we compared the apoptotic response of wild-type and *Drp1*^*−/−*^ ESCs to ER stress or oxidative stress. We observed that a significantly higher proportion of *Drp1*^*−/−*^ cells treated with thapsigargin or sodium arsenite show MOMP (Figure 5C). We next analysed the kinetics of MOMP in response to BH3 peptides and observed that both BIM and BID BH3 peptides were more efficient at inducing mitochondrial outer membrane depolarization in *Drp1*^*−/−*^ ESCs than wild-type cells (Figure 5E). This indicates that loss of *Drp1* is sufficient to lower the apoptotic threshold of ESCs (Figure 2E). Importantly, this change in sensitivity to cell death induction was unlikely to be a secondary consequence of a disruption of the metabolism of mutant cells, as *Drp1*^*−/−*^ ESCs did not show decreased glycolysis or oxidative phosphorylation rates when compared to wild-type cells. Instead, mutant cells displayed an increased spare respiratory capacity, an increased response to pyruvate and an increased glycolytic rate (Supplementary Figure 3B-E), suggesting that mitochondrial fission enhances the bioenergetic rate of pluripotent cells.

DRP1 has been shown to be involved in the remodelling of the mitochondrial cristae and the subsequent release of cytochrome C into the cytoplasm^17^, ^18^, an event downstream of MOMP. We therefore analysed if cytochrome C release is compromised in *Drp1*^*−/−*^ ESCs. We observed that *Drp1*^*−/−*^ ESCs displayed lower levels of cytoplasmic cytochrome C upon thapsigargin treatment (Supplementary Figure 3F). This suggests that DRP1 is likely to play at least two roles in the apoptotic response in pluripotent cells. First, it impedes the pro-apoptotic roles of BCL-2 family members and second it promotes cytochrome C release by helping the remodelling of the mitochondrial cristae.

Given the roles of DRP1 in determining the apoptotic response of ESCs, we next wanted to analyse its importance during ESCs differentiation. During differentiation p-DRP1 levels decrease (Figure 4C) and therefore we analysed the effects of *Drp1* overexpression (*Drp1*^*OE*^) in differentiating ESCs. We have previously shown that culturing ESCs for 3 days in N2B27 leads to a post-implantation epiblast-like state^32^. We therefore cultured *Drp1*^*OE*^ cells for 3 days in N2B27 and found that this led to sustained DRP1 expression and increase in p-DRP1 levels (Supplementary Figure 4A-C and Supplementary Figure 5A-D). Importantly, this increase in p-DRP1 did not prevent exit of naïve pluripotency but induced mitochondria to adopt a more rounded morphology (Figure 6A-C and Supplementary Figure 5A-B). We therefore tested the effect on the apoptotic response of a sustained increase in p-DRP1 during the onset of ESC differentiation. We found that after 3 days in differentiation conditions, *Drp1*^*OE*^ cells displayed not only lower basal cleaved caspase 3 levels (Figure 6D and Supplementary Figure 5A-C), but also significantly reduced levels of cleaved Caspase 3 in response to treatment with thapsigargin, sodium arsenite or ABT-737 (Figure 6E-G and Supplementary Figure 5A,C-D). Together with our findings in *Drp1*^*−/−*^ ESCs, these results suggest that the decrease in DRP1 and p-DRP1 levels observed during exit of pluripotency play an important role in sensitising cells to apoptosis.

**Figure 6.**
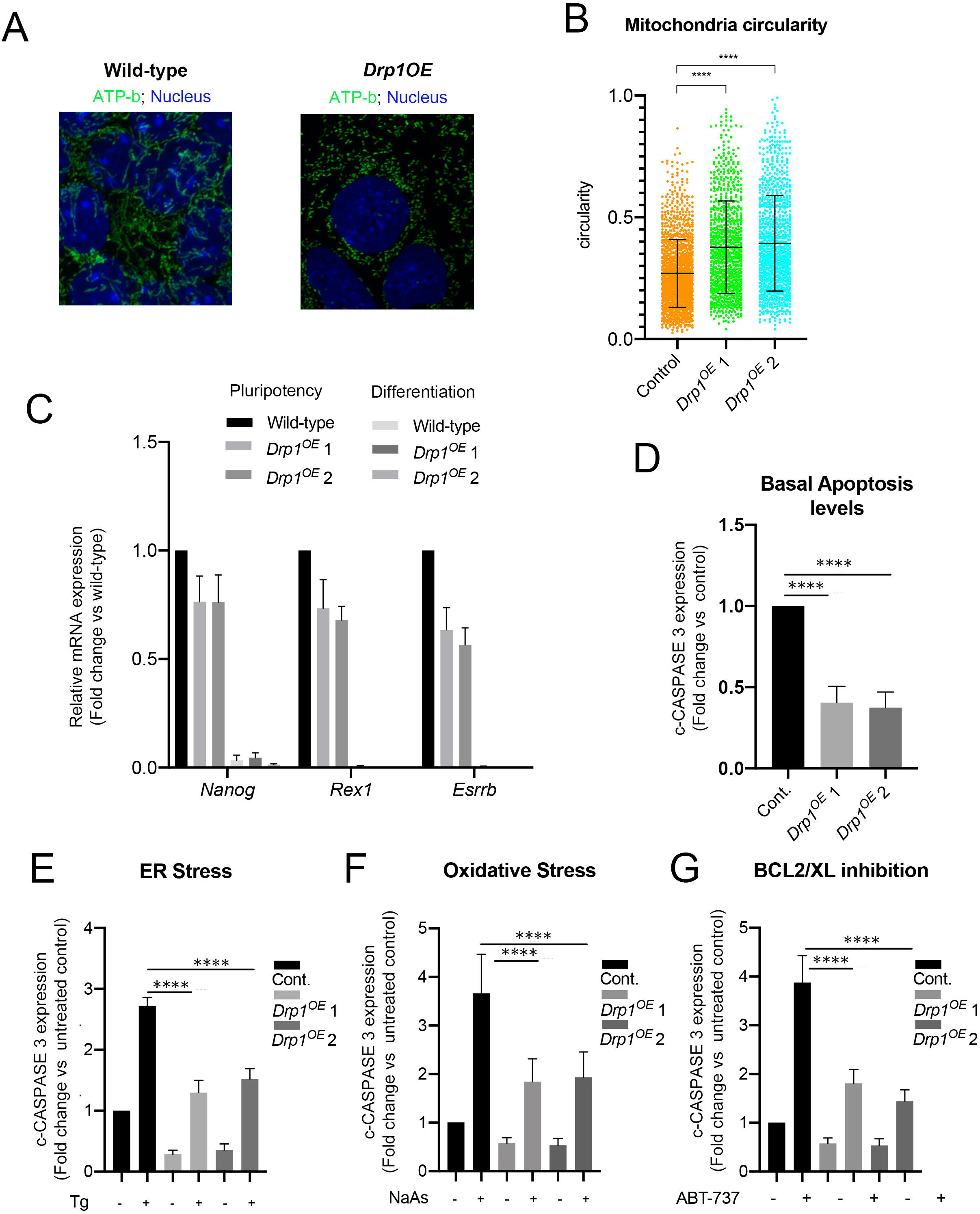
DRP1 over-expression inhibits the apoptotic response during the onset of pluripotent stem cell differentiation. **A.** ATP-b immunostaining showing mitochondrial morphology in wild-type and *Drp1*^*OE*^ cells during differentiation. **B.** Circularity measurement of mitochondrial particles from ATP-b immunostained images of wild-type and *Drp1*^*OE*^ cells at day 3 of differentiation in N2B27. **C.** Quantitative RT-PCR showing gene expression levels of naïve and primed pluripotency markers in wild-type and *Drp1*^*OE*^ ESCs and at day 3 of differentiation in N2B27. Gene expression normalized against *Gapdh*. **D.** Fold change in basal cleaved-CASPASE 3 levels in wild-type and *Drp1*^*OE*^ cells at day 3 of differentiation in N2B27. **E.** Fold change in cleaved-CASPASE 3 levels in wild-type and *Drp1*^*OE*^ cells untreated or treated with 1µM Thapsigargin for 5h. **F.** Fold change in cleaved-CASPASE 3 levels in wild-type and *Drp1*^*OE*^ cells untreated or treated with 1µM sodium arsenite for 5h. **G.** Fold change in cleaved-CASPASE 3 levels in wild type and *Drp1*^*OE*^ cells untreated or treated with 1µM ABT-737 for 5h at day 3 of differentiation in N2B27. Protein levels in **D. E. F.** and **G.** are normalized against α-TUBULIN and graphs show protein expression levels relative to wild-type cells. Average of 3 (C, E, F) or 5 (D) independent experiments +/-SEM is shown. Students T-Test (D) or 2-way ANOVA with Šidák correction (E, F, G) ****p<0.0001. The statistical comparisons are made to untreated cells.

## Discussion

The removal of aberrant cells during development is important to prevent them from contributing to further development and the germline. To facilitate this elimination cells become hypersensitive to apoptosis during the onset of differentiation^6^, ^11^, ^33^, ^34^. Here we show that DRP1 regulates apoptotic priming in these cells via influencing mitochondrial dynamics. We show that upon exit of naïve pluripotency mitochondria form complex networks due to a decrease in mitochondrial fission induced by a loss of DRP1 phosphorylation. We also demonstrate that this loss of DRP1 activity changes the apoptotic threshold, as deletion of *Drp1* facilitates the early stages of apoptosis and its over-expression protects against apoptosis. Together, these findings highlight the pivotal role that DRP1 plays in modulating the response to cell death during differentiation.

When considering how DRP1 levels might affect the apoptotic response, a number of possibilities arise. In the first instance, it has been shown that mitochondrial shape affects the ability of BAX to permeabilise the outer mitochondria membrane. It was found that mitochondrial fusion establishes a mitochondrial size that is permissive to the function of pro-apoptotic BCL2 family members and that mitochondrial hyper-fragmentation inhibited the ability of BAX to associate with and permeabilise the outer mitochondria membrane^23^. In line with this possibility, the rounded and fragmented shape of the mitochondria of naive pluripotent cells due to the high levels of fission induced by DRP1 would help explain the high apoptotic threshold of these cells.

An alternative, but not mutually exclusive, possibility that could explain how DRP1 affects the apoptotic response, is by modulating the entry of calcium into the mitochondria. It is well known that the degree of contact between the endoplasmic reticulum (ER) and mitochondria has profound implications for the function of each of these organelles^13^, ^35^. For example entry of calcium into the mitochondria is thought to be regulated by ER-mitochondria contacts. Moreover, ER-mitochondria contact sites are thought to be a site of reactive oxygen species (ROS) signalling, which also affects calcium signals delivery to the mitochondria^36^. We observe that both mutation of DRP1 or its overexpression changes the apoptotic response to thapsigargin, a drug that increases calcium levels in the cytoplasm, and to sodium arsenite, which induces oxidative stress. Therefore, our results raise the possibility that by regulating mitochondrial shape, DRP1 levels control the degree of contact between the ER and mitochondria, with the formation of elongated mitochondrial networks induced by decreased fission during differentiation favouring increased mitochondria ER-contacts and in this way facilitating calcium entry into the mitochondria.

It is worth highlighting that the involvement of DRP1 in the early apoptotic events involving permeabilization of the mitochondrial membrane are separable from the downstream release of cytochrome C, which others have shown and we also find is promoted by DRP1^17^, ^18^, ^37^. In principle, this would represent two opposing roles for DRP1: slowing mitochondrial membrane permeabilization, but facilitating cytochrome C release. One way to reconcile these potentially opposing roles is that either each one requires different levels of DRP1, with the release cytochrome C being possible with just baseline DRP1 levels. Alternatively, different post-translational modifications of DRP1 may lead to altered function. Indeed, DRP1 has been shown to be phosphorylated at 3 different sites, as well as being S-nitrolisilated, O-GlcNAcylated, SUMOylated and ubiquitinated^13^, ^38^ and each of these modifications are associated with different roles of DRP1. We have only analysed S616 phosphorylation and found it to be lost during ESC differentiation, but to gain a more in depth understanding of the roles of DRP1 in the apoptotic response, it would be important to analyse how a broader range of post-transcriptional modifications differ between the naïve and primed pluripotent states.

Our results contrast with the roles that others have identified for mitochondrial dynamics in pluripotency. For example over-expression of *Mff*, a fission regulator, inhibits the expression of neural markers during pluripotent stem cell differentiation^39^, suggesting that inhibiting mitochondrial elongation disrupts differentiation. Similarly, mutation of *Mtch2*, a potential fusion regulator, delays exit from naïve pluripotency^40^, leading to the suggestion that mitochondrial fusion promotes differentiation. In contrast to this we find that *Drp1* null ESCs, which have hyper-fused mitochondria, show normal pluripotency gene expression and over-expression of *Drp1* does not affect exit from naïve pluripotency. What these results suggest, is that mitochondrial elongation *per se* is not linked to exit of pluripotency, but rather it is likely that other mitochondrial processes regulated by *Mff* and *Mtch2* have an impact on the onset of differentiation.

Finally, our work has implications that transcend early mammalian development as DRP1 has been shown to play roles in tumour progression. For example, in glioblastoma DRP1 activation is correlated with poor prognosis. Mechanistically this is explained as brain tumour initiating cells have fragmented mitochondria and require high p-DRP1 levels for their survival^41^. Similarly, in pancreatic ductal adenocarcinomas, oncogenic *Ras* mutations induces mitochondrial fragmentation. Reversion of this phenotype by knock-down of *Drp1* inhibits tumour growth^42^, ^43^, further highlighting the potential importance of DRP1 for tumour progression. Our findings that in embryonic stem cells DRP1 promotes cell survival, raises the possibility that part of this oncogenic role is through the regulation of the apoptotic response. Understanding the players downstream of DRP1 will therefore likely open new avenues for our understanding of transformation.

## Acknowledgements

We would like to Massimo Signore and Juan Pedro Martinez Barbera for critical discussion. We thank Stephen Rothery for guidance and advice with confocal microscopy. Gratitude also goes to James Elliot for performing cell sorting. Research in the Tristan Rodriguez lab was supported by an MRC project grant (MR/N009371/1) and by the British Heart Foundation centre for research excellence. Barbara Pernaute was supported by an EMBO Long Term Fellowship (1340-2010) and a Marie Curie Intra-European Fellowship (FP7-PEOPLE-2010-IEF n° 273884). Salvador Perez Montero has been a recipient of an EMBO long-term Fellowship (846-2015) and a Marie Curie Intra-European Fellowship (H2020-MSCA-IF-2015 nº 709010). Work in the Meier lab is funded by Breast Cancer Now (CTR-QR14-007). Guy A. Rutter was supported by Wellcome Trust Senior Investigator (WT098424AIA) and Investigator (212625/Z/18/Z) Awards, MRC Programme grants (MR/R022259/1, MR/J0003042/1, MR/L020149/1) and Experimental Challenge Grant (DIVA, MR/L02036X/1), MRC (MR/N00275X/1), and Diabetes UK (BDA/11/0004210, BDA/15/0005275, BDA 16/0005485) grants. This project has received funding from the European Union via the Innovative Medicines Initiative 2 Joint Undertaking under grant agreement No 115881 (RHAPSODY) to Guy A. Rutter. We acknowledge NHS funding to the NIHR Biomedical Research Centre. Alejandra Tomás is funded by the MRC project grant MR/R010676/1.

## Materials and Methods

### Stem Cell culture and treatments

All cells were cultured at 37°C in an atmosphere with 5% CO_2_. Reagents used for tissue culture were obtained from Invitrogen unless otherwise stated. Mouse embryonic stem cells (ESCs) were cultured on 0.1% gelatin-coated flasks (Nunc, Thermo Fisher) in GMEM containing with 10% (v/v) foetal calf serum (FCS; Seralab), 1X non-essential amino acids, 2 mM L-glutamine, 0.1 mM β-mercaptoethanol and supplemented with homemade leukaemia inhibitory factor (1:500, LIF). ESCs were routinely dissociated with trypsin and cryopreserved in 10%DMSO in FCS.

Epiblast Stem Cells (EpiSCs) were cultured on FCS coated dishes in N2B27 medium (100mL DMEM F12, 100mL Neurobasal, 1mL N2, 2mL B27 without retinoic acid, 2mM L-Glutamine, 50mM-mercaptoethanol) containing 20µg/ml Activin A (R&D Systems) and 12ng/ml bFGF (R&D Systems). Cells were passaged by mechanical disruption as previously described^44^. E14 EpiSC were derived from E14 mESCs as previously described^45^. All experiments were performed using cells in passage between 20 and 30.

To induce ESCs differentiation cells were seeded onto plates coated with fibronectin (Merk) and cultured in N2B27 media (Neurobasal media; DMEM F12 media, 0.5 x B27 supplement; 0.5 × N2 supplement; 0.1mM 2-mercaptoetanol; 2mM glutamine; all Thermo Fisher Scientific) during 3 days to allow differentiation.

To induce *Dicer* deletion *Dicer*^*fx/fx*^ ESCs and EpiSCs^11^, ^46^ were cultured in the presence of 0.3 mM 4-OH-Tamoxifen for three days and left untreated from the third day onwards as previously described^11^.

For BCL2/BCL-XL inhibition ABT-737 (Selleckchem) was added to the media for 24 h at the stated concentration. Oxidative stress and ER stress were induced by adding sodium arsenite (Sigma Aldrich) or Thapsigargin (Sigma Aldrich) for 16 h at the stated concentrations. To induce *Bim* knockdown, a previously tested *Bim* siRNA (Mm_Bcl2l11_2 FlexiTube siRNA, Qiagen) ^11^ was transfected into EpiSCs at a final concentration of 75nM using HiPerFect transfection reagent (Qiagen) according to manufacturers’ instructions. Transfection of Flexi-Tube Negative Control siRNA (Qiagen) at a final concentration of 75nM was used as negative control.

### Flow cytometry analysis: Annexin V staining and MMP measurement

For apoptosis detection by flow cytometry, Annexin V-APC (Thermo Fisher Scientific) was used in combination with Propidium Iodide (Sigma) according to manufacturer’s instructions. Briefly, approximately 2×10^5^ cells were stained in 100µl of Annexin V Binding Buffer (0.1% BSA in 10 mM HEPES, 140 mM NaCl, 2.5 mM CaCl2, pH7.4) containing APC conjugated Annexin V for 15 minutes in the dark, after which 0.1 mg/ml propidium iodide was added and the samples immediately analysed by flow cytometry. Loss of mitochondrial membrane potential was measured using the fluorescent dyes DiOC6 or TMRM. Briefly, 2×10^5^ cells were re-suspended in PBS containing 40 nM DiOC6 (Sigma) or 100nM TMRM (T668, ThermoFisher Scientific), incubated for 15 min at 37°C and analyzed by flow cytometry. Data was acquired with a BD LSRII cytometer and analyzed with the FlowJo software (BD).

### Stem cells immunofluorescence

For immunostaining ESCs and EpiSCs were fixed for 10min in 4%PFA at room temperature, permeabilised in 0,4% Triton-X100/PBS for 5 minutes at room temperature, blocked in 10%BSA/ 0,1% Triton X-100/PBS and incubated overnight at 4°C in primary antibody diluted in 1%BSA/0,1%Triton X-100 (anti-Cleaved Caspase 3 Asp175, Cell Signalling, 1/100; anti-Nanog (14-5761□80 eBioscience, 1/100, anti ATP-b (Ab14730, Abcam – 1:200). Alexa-Fluor conjugated secondary antibodies (Thermo Fisher Scientific) were used at 1/500 dilution in 1%BSA/0,1%Triton X-100. Cells were mounted for visualization in Vectashield with DAPI (Vector Laboratories). Images were acquired with a Zeiss confocal microscope and analysed with the Fiji software^47^.

#### Mitochondrial Staining

Cells were washed with PBS and fixed with 3.7% formaldehyde diluted in serum free media for 15 min at 37°C, 5% CO_2_. Cell were washed two times with PBS and incubated with pre-cooled acetone for 5 min at −20°C. Cells were washed two times with PBS and incubated with blocking/permeabilization (5% BSA, 0.4% Triton-X in PBS) solution for 30 min at RT before incubating with the primary antibody O/N at 4°C. Excess antibody was removed and cells washed three times in PBS, then incubated with the secondary antibody for 45 min at RT. Before mounting with Vectashield with DAPI, secondary antibody was removed and cell washed again three times in PBS.

Images were acquired in a LSM Z800 Confocal microscope and processed with FIJI. Images for deconvolution were acquired with the same microscope and further processed with the software Huygens. Deconvolution analysis was performed with the support and advice from Mr. Stephen Rothery from the Facility for Imaging by Light Microscopy (FILM) at Imperial College London.

Primary antibodies used for immunofluorescence: Tom20 (1/100, Santa Cruz), ATP-b (1:100, Abcam), Nanog (1:100, eBioscience), Gata4 (1:100, Santa Cruz), Sox2 (1:100, R&D). Alexa-Fluor conjugated secondary antibodies (Invitrogen) were used at a concentration of 1:600.

Mitochondria circularity measurements were done with a plugin from FIJI that calculates object circularity according to the formula circularity=4π(area/perimeter^2^). A circularity value of 1.0 indicates a perfect circle. As the value approaches 0.0, it indicates an increasingly elongated polygon. The calculations were done on ATP-b immunostained images.

### Western blot analysis

Western blot was performed according to established protocols described elsewhere^11^. Briefly, protein samples were collected in Laemmli buffer and denatured for 10 minutes at 95°C. All samples were run in polyacrylamide gels and transferred to nitrocellulose membranes. Blocking was performed in 5% milk in TBST buffer and primary antibody incubation was done overnight at 4°C in TBST containing 5% BSA. The following antibodies were used at the stated concentration: anti-Bim/Bod (Enzo, 1/1000), anti-Puma (Abcam, 1/1000), anti-Bid (Cell Signaling, 1/1000), anti-Bad (Santa Cruz Biotechnology, 1/500), anti-Bax N20 (Santa Cruz Biotechnology, 1/1000), anti-Bcl2 (BioLegend, 1/500), anti-Bcl-XL (Santa Cruz Biotechnology, 1/1000), anti-MCL1 (Rockland, 1/10000), anti-Bcl2A1 (R&D Systems, 1/500), anti-Hexokinase II (Cell Signaling, 1/1000), anti-ATP-b (Abcam, 1/1000), anti-c C (BD Pharmigen, 1/1000), anti-Erk1/2 (Sigma, 1/20000), anti-Cleaved Caspase 3 Asp175 (Cell Signaling, 1/1000), anti-alpha Tubulin (Cell Signaling, 1/2000), anti-beta Actin (Santa Cruz Biotechnology, 1/1000), anti-Bak (Millipore, 1/1000) anti-Oct3/4 (Santa Cruz Biotechnology, 1/1000), anti-Nanog (eBiosciences, 1/1000), anti-Tom20 (Santa Cruz Biotechnologies, 1/1000), anti-Drp1 (Cell Signalling, 1/1000) anti-pDRP1 (S616) (Cell Signaling 1/1000), anti-Mfn1 (Abcam, 1/1000), anti-Mfn2 (Abcam, 1/1000), anti-Opa1 (BD, 1/1000). Western blot quantification was performed using the Fiji software^47^.

#### Mitochondria purification and cytochrome C release assay

For mitochondria purification cells were trypsinized and washed twice in 10 packed cell volumes of 1mM TrisHCl pH 7.4, 0.13M NaCl, 5mM KCl, 7.5 mM MgCl_2_ followed by centrifugation at 370g for 10 minutes. After the second wash pelleted cells were re-suspended in 6 packed cell volumes of homogenization buffer (10mM TrisHCl pH6.7, 10mM KCl, 0.15 mM MgCl_2_, 1mM PMSF, 1mM DTT) and incubated for 10 minutes on ice. Cells were homogenized in a glass homogenizer until achieving approximately 60% cell breakage. Homogenate was poured into a tube containing 1 packed cell volume of 2M sucrose solution, mixed and centrifuged at 1200g for 5 minutes to pellet unbroken cells, nuclei and debris. This treatment was repeated twice discarding the pellet, followed by 10 minutes centrifugation at 7000g in order to pellet the mitochondria. Mitochondrial pellet was re-suspended in 3 packed cell volumes of mitochondrial suspension buffer (10mM TrisHCl pH6.7, 0.15mM MgCl_2_, 0.25% sucrose, 1mM PMSF, 1mM DTT) and centrifuged for 5 minutes at 9500g. Final mitochondrial pellet was re-suspended in 1x Laemmli Buffer and boiled at 95ºC for 10 minutes for western blot analysis.

For the separation of cytosolic and membrane fractions in order to investigate cytochrome C release, cells were washed twice with ice cold PBS and incubated for 10 minutes rocking on ice in Digitonin Buffer (20mM HEPES/KOH pH7.5, 100mM sucrose, 2.5mM MgCl_2_, 100mM KCL, 1mM DTT, 0.0025% digitonin) supplemented with Complete protease inhibitors (Roche). The supernatant was collected as cytosolic fraction and Triton X-100 Extraction Buffer^48^ was added to the plates followed by 30 minute incubation rocking on ice. The resulting supernatant was taken as membrane fraction.

### Embryo culture, treatment and immunofluorescence

E3.5 embryos were obtained by flushing of the uterus at 3.5 days post coitum and cultured in M16 media containing DMSO, 1µM Etoposide (Sigma) or 2µM ABT-737 (Selleckchem). E6.5 embryos were dissected from pregnant females at 6.5 days post coitum and cultured in N2B27 media (see Epiblast Stem Cell culture conditions) containing DMSO, 1µM Etoposide or 2µM ABT-737.

For immunostaining, embryos were fixed in 4% PFA/0.1%Tween/0.01% Triton X-100/PBS (10 min for 3.5dpc and 4.5dpc, 20min for 6.5dpc) at room temperature, permeabilized in 0.5% Triton X-100/PBS for 15min (3.5dpc and 4.5dpc) or 20min (6.5dpc), washed in 0.1% Triton X-100/PBS and blocked in 2% Horse serum in 0.1% Triton X-100/PBS for 45 minutes at room temperature. Primary antibodies (anti-Cleaved Caspase 3 Asp175, Cell Signaling, 1/100; anti-p53 (1C12), Cell Signalling, 1/100, anti ATP-b, (Ab14730, Abcam – 1:200) were incubated overnight at 4°C in 0.2% Horse serum in 0.01% Triton X-100/PBS. Embryos were incubated with Alexa-Fluor conjugated secondary antibodies (Thermo Fisher Scientific) for 1h at 4°C and counterstained with 6ug/ml Hoechst for nuclear visualization. Images were acquired with a Zeiss confocal microscope and analyzed with the Fiji softwar^47^.

#### Embryo image quantification

Corrected cell fluorescence was measured using (Fiji, Image J) as previously described (reference below). An outline was drawn around each embryo. Area, mean fluorescence and integrated density were measured. In addition, adjacent areas were also selected an measured as background readings. Corrected cellular fluorescence (CCF) was calculated using the formula: CCF = integrated density – (area of selected cell × mean fluorescence of background readings. Cleaved Caspase 3 signal was normalised to DAPI^49^.

### BH3 profiling

The assay was performed following the protocol described by Anthony Letai’s laboratory^27^. Briefly, 15μL of 2x peptides diluted in DTEB buffer (135mM threalose, 10mM HEPES KOH pH 7.5, 50mM KCl, 20μM EGTA, 20μM EDTA, 0.10% BSA, 5mM succinic acid) were added to each well of a dark 384 well plate (Nunc). Cells were dissociated with trypsin, counted and resuspended at 2.67×10^6^ cells/mL. Equal volume of cells and dying mix (4x digitonin (0.01%), 4x JC-1 (4mM) and Oligomycin (40μg/mL)) were incubated for 10 minutes at room temperature protected from light. 15μL of this mix was added per well of the 384 well plate. DMSO and CCCP (Sigma) were used as no-depolarization and full depolarization controls. BID, BIM, BMF and a peptide control were tested in this profile. Each sample was loaded by triplicate and at least three biological replicates were analysed. Fluorescence was measured every 5 minutes for a period of 70 minutes, with a 544/590 filter in a FluoStar Omega plate reader (BMG Omega). Percentage of depolarization was calculated by normalizing the data to the membrane potential of cells that have not been exposed to peptides but have been treated with DMSO (vehicle) or CCCP, a protonophore that causes complete mitochondrial depolarization^27^.

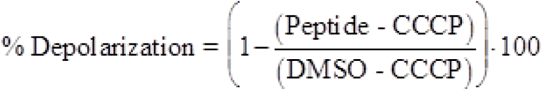

| PEPTIDE | SEQUENCE |
| --- | --- |
| **BIM** | Ac-MRPEIWIAQELRRIGDEFNA-NH<sub>2</sub> |
| **BID** | Ac-EDIIRNIARHLAQVGDSMDRY-NH<sub>2</sub> |
| **BFM** | Ac-HQAEVQIARKLQLIADQFHRY-NH<sub>2</sub> |
| **CONTROL** | Ac-EQWAREIGAQARRMAADLNA-NH<sub>2</sub> |

### Generation of *Drp1* Knock-Out and *Drp1* Overexpressing cells

*Drp1* knockout ESCs were generated by CRISPR-Cas9 mediated deletion of *Drp1* exon 2. sgRNA guides flanking *Drp1* exon2 were cloned into the PX459 vector (Addgene) ^50^: *Drp1* exon 2 upstream sgRNA: 5’ TGGAACGGTCACAGCTGCAC 3’; *Drp1* exon 2 downstream sgRNA: 5’ TGGTCGCTGAGTTTGAGGCC 3’. E14 ESCs were co-transfected with 1ug of each sgRNA expression using Lipofectamine 2000 (Invitrogen) according to manufacturer’s instructions. As control E14 ESCs were transfected in parallel with equal amount of empty PX459 plasmid. Following 6 days of Puromycin selection, single colonies were picked from both *Drp1* sgRNA and empty vector transfected ESCs and screened for mutations. *Drp1* exon 2 deletion was confirmed by PCR genotyping using the following primers: *Drp1*_genot F: 5’ GGATACCCCAAGATTTCTGGA 3’; *Drp1*_genot R: 5’ AGTCAGGTAATCGGGAGGAAA 3’, followed by Sanger Sequencing.

*Drp1* overexpressing cells were generated by transfecting ESCs with a pCAG-*Drp1* plasmid. To generate this plasmid the *Drp1* cDNA (Addgene plasmid 45160) was cloned into a pCAG mammalian expression plasmid. E14 ESCs were transfected with 2 ug of the pCAG-*Drp1* plasmid using Lipofectamine 2000 (Invitrogen) according to manufacturer’s instructions. Following 6 days of Puromycin selection, single colonies were picked and analysed for DRP1 expression by western blot.

### RNA Extraction and Quantitative RT-PCR

RNA was extracted with the RNeasy mini kit (Qiagen) and SuperScript III reverse transcriptase (Thermo Fisher Scientific) was used for cDNA synthesis according to manufacturer’s instructions. Quantitative RT-PCR was performed by amplification with Lightcycler 480 SYBR green Master (Roche). The primers used are listed in Table 1. RNA samples from wild type and mutant clones were collected from 3 independent experiments.

**Table 1:**
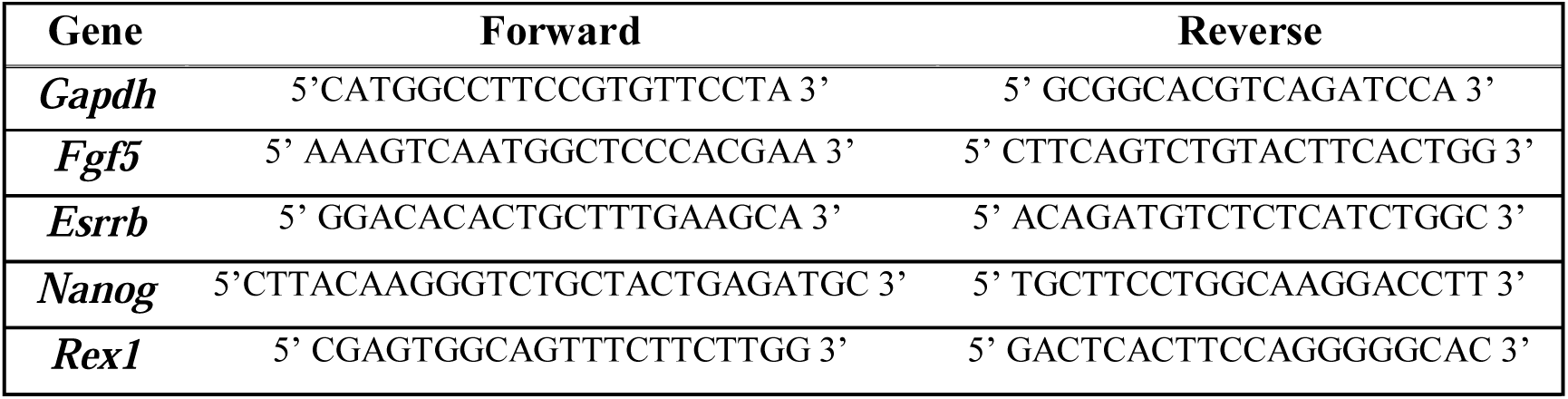
Primers used in quantitative RT-PCR

For miRNA q-PCR total RNA was extracted using the *mir*Vana miRNA isolation kit (Ambion) and cDNA was synthesized with the TaqMan miRNA reverse transcription kit (Applied Biosystems). qPCR for individual miRNAs was performed using TaqMan probes and TaqMan universal PCR master mix (Applied Biosystems). miRNA expression was normalized against sno135.

### Seahorse Analysis

The metabolic function of cells was assessed by extracellular flux analysis using Seahorse XF24 (Agilent Technologies, UK). On the day prior to the assay, 5×10^4^ cells per well were seeded on 0.1% gelatin-coated (Sigma, UK) 24-well plates and grown in 300 µL of pluripotency maintenance conditions. Cells were washed, just before the assay, with the assay media and left with a final volume of the 600 µL per well. The plate was then equilibrated on a non-CO_2_ incubator at 37ºC for 30 min. The assay media consisted in unbuffered DMEM (D5030 – Sigma, UK), reconstituted with 1.83 g.L^−1^ of NaCl in dH_2_O, that was supplemented on the day of the assay according to the test performed. For the OCR measurements the assay media was supplemented with 0.5 g.L^−1^ of glucose (Sigma, UK) and 2 mM of L-glutamine (Life Technologies, UK), while for the ECAR measurements the media was supplemented with 1 mM of Sodium Pyruvate and 2 mM of L-glutamine (both from Life Technologies, UK), pH 7.4 at 37ºC.

Assays were performed with 5 biological replicates of each cell type and 4 wells were left without cells for background correction measurements. Both ECAR and OCR measurements were performed on the same plate. The assay parameters for both tests were calculated following the Seahorse assay report generator (Agilent Technologies, UK).

The protocol for the assay consisted of 4 baseline measurements and 3 measurements after each compound addition. Compounds (all from Sigma, UK) used in OCR and ECAR tests were prepared in the supplemented assay media. For the OCR, test the following compounds were added: 2.5 µM oligomycin (OM), 300 nM Carbonyl cyanide-4-(trifluoromethoxy) phenylhydrazone (FCCP) and a mixture of rotenone and antimycin A at 6 µM each (R&A). For the ECAR test, the following compounds were added: 2.5 mM and 10 mM of glucose, 2.5 µM of oligomycin (OM), and a 50 mM of 2-deoxyglucose (2-DG).

At the end of the assay, cells were fixed and stained with Hoechst. Both OCR and ECAR were normalised to cell number, determined by manual cell counts using FIJI software.

## Statistical methods

Statistical analysis was performed using GraphPad Prism software. Statistical methods used are indicated in the relevant figure legends. No randomization or blinding was used in experiments. Sample sizes were selected based on the observed effects and listed in the figure legends.

---

**Supplementary Figure 1.**
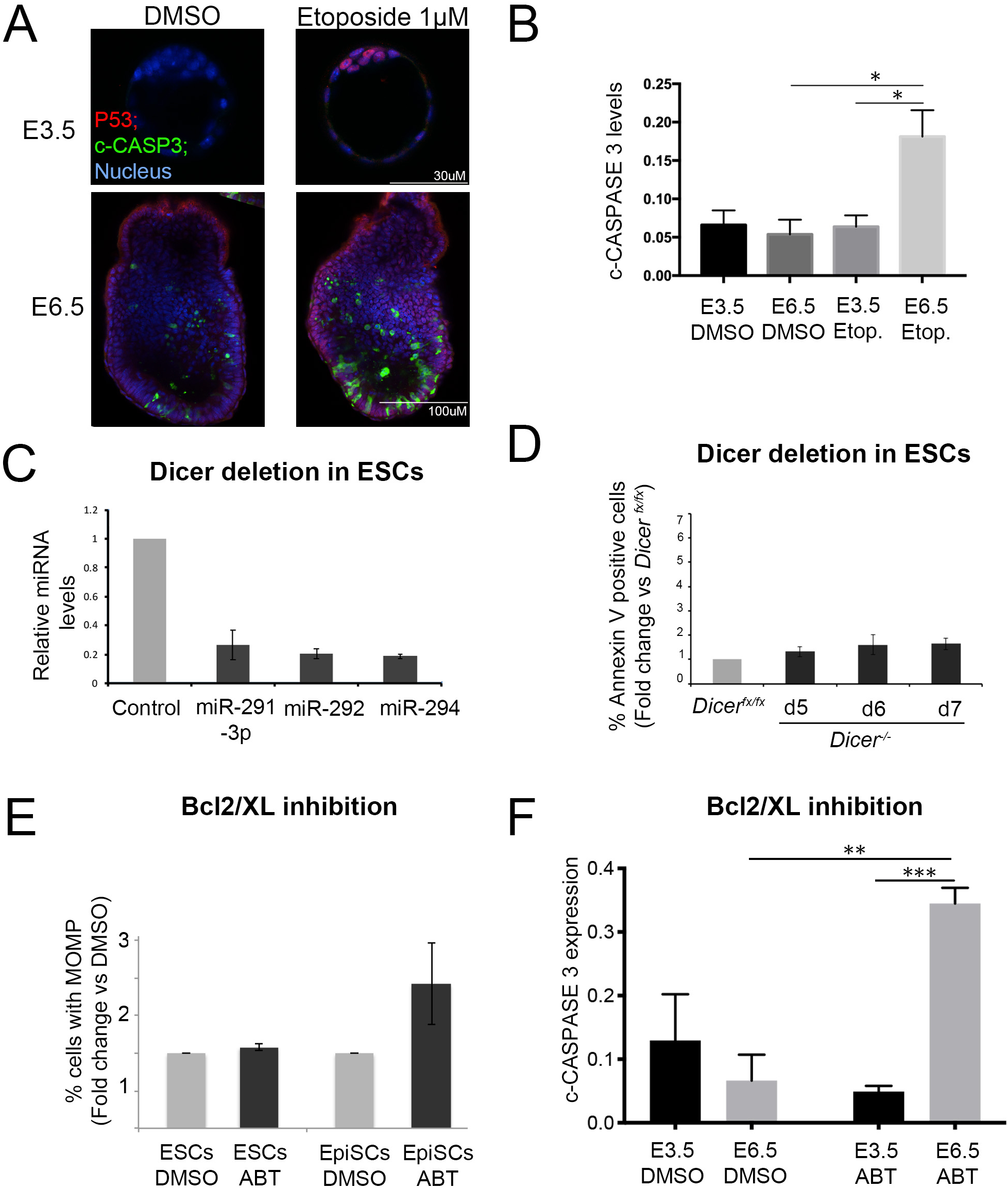
Enhanced sensitivity to apoptosis in primed pluripotent stem cells. **A.** Levels of apoptosis as measured by cleaved-caspase 3 in E3.5 and E6.5 embryos cultured for 1.5h in the presence of DMSO or 1µM Etoposide. **B**. Quantification of cleaved-caspase 3 staining in A. Dmso: E3.5 (n= 3); E6.5 (n=4); 1µM Etoposide: E3.5 (n= 6); E6.5 (n=10). **C.** Quantitative RT-PCR showing change in miRNA relative expression 6 days after *Dicer* deletion in ESCs. Expression normalized against sno135. Fold change vs un-deleted cells is shown. **D.** Change in % of Annexin V positive ESCs at different time points after *Dicer* deletion relative to un-deleted cells. **E**. Change in % of ESCs and EpiSCs with loss of mitochondrial membrane potential (MOMP) after 24h treatment with 5µM ABT-737, relative to dmso treated cells. **F.** Cleaved-CASPASE 3 levels in E3.5 and E6.5 embryos treated with dmso or 2µM ABT-737 for 1.5h. E3.5 dmso (n=5), ABT (n=5); E6.5 dmso (n=6) ABT (n=6). Average of a minimum of (B,F) 3, (D) 4 or (E) 5 experiments +/-SEM is shown. 2-way ANOVA with a Turkey correction (B, G) *p<0.05, **p<0.01 or ***p<0.001.

**Supplementary Figure 2.**
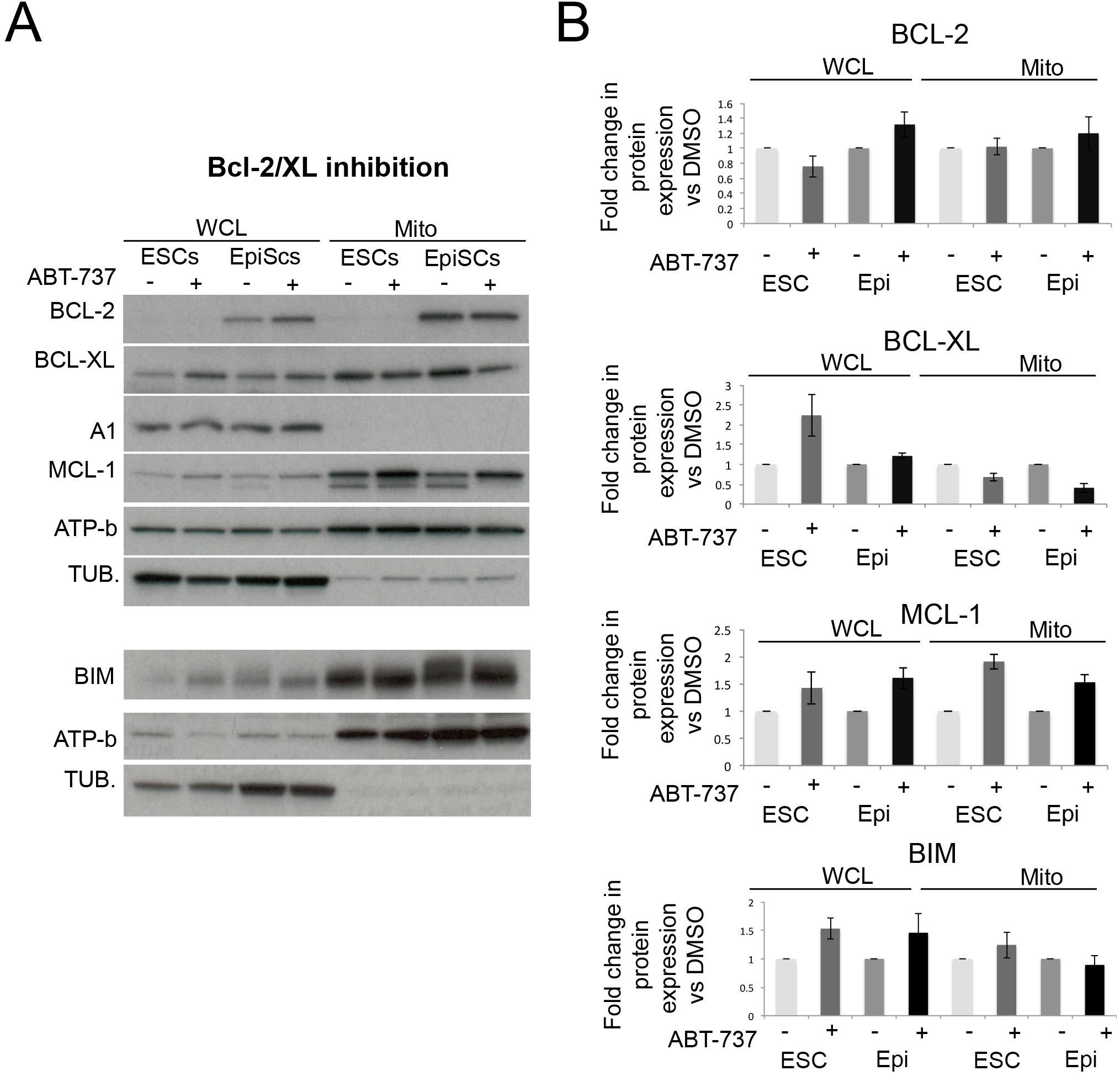
Expression of BCL-2 family members upon BCL-2/XL inhibition. **A.** Levels of anti-apoptotic factors and BIM in whole cell extracts and mitochondrial extracts from ESCs and EpiSCs treated with dmso or 5µM ABT-737 for 24h. A representative western blot is shown. **B**. Levels of anti-apoptotic proteins and BIM in whole cell and mitochondrial extracts from ESCs and EpiSC (Epi)s treated with dmso or 5µM ABT-737 for 24h. Protein level is normalized against α-TUBULIN (TUB.) in whole cell extracts and against ATP-b in mitochondrial extracts. Fold change of ABT versus dmso treated is shown. Average of 4 experiments +/-SEM is shown.

**Supplementary Figure 3.**
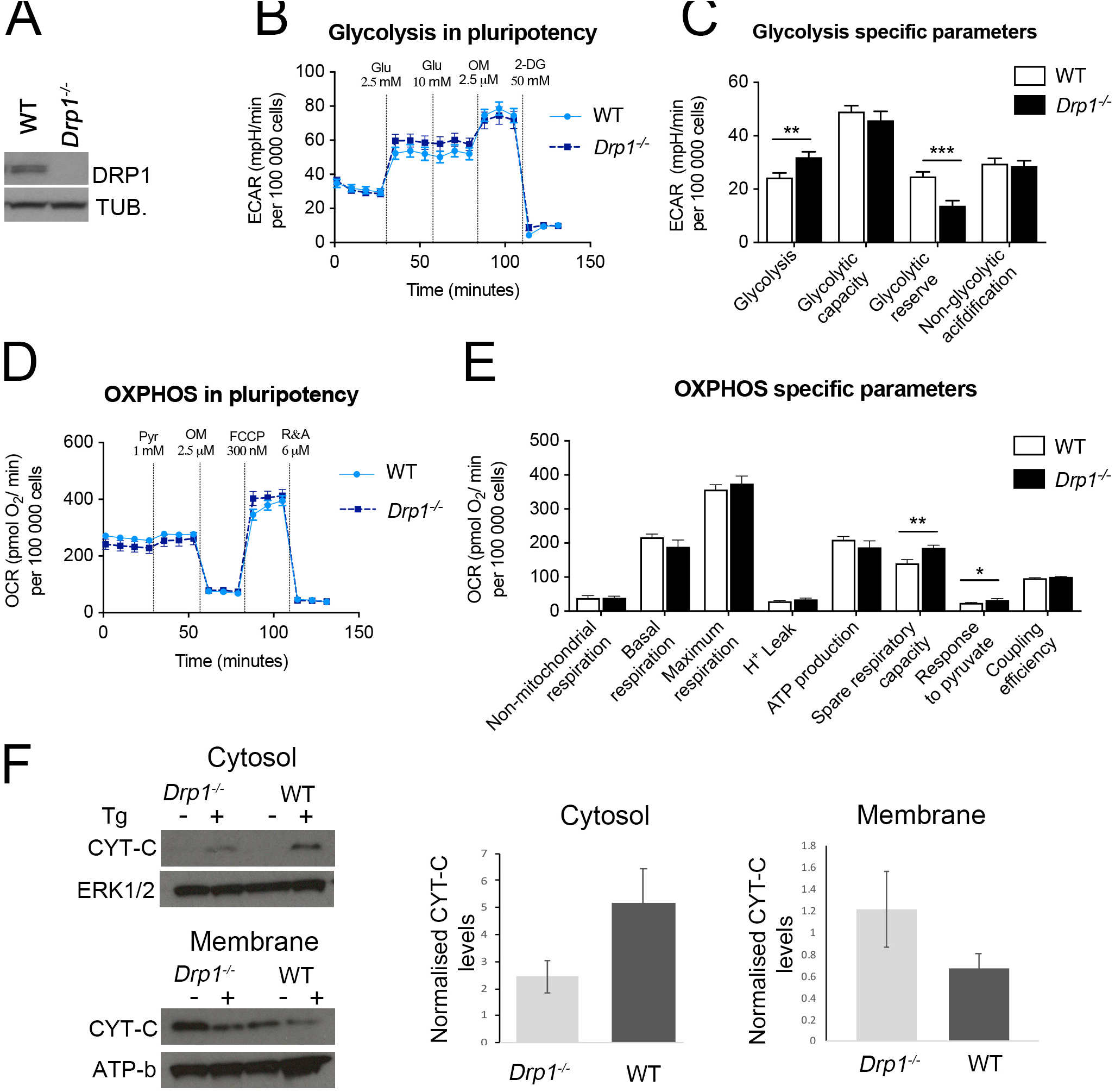
Metabolic profile of wild-type and *Drp1*^*−/−*^ ESCs. **A.** Total DRP1 protein levels in wild type and *Drp1*^*−/−*^ ESCs. **B.** Extracellular acidification rate (ECAR) during the glycolysis stress test. **C** Metabolic parameters assessed during a glycolysis stress test. **D.** Oxygen consumption rate (OCR) during the mitochondria stress test. **E.** Metabolic parameters assessed after the mitochondria stress test. **F.** CYTOCHROME C (CYT-C) protein levels in cytosolic and membrane fractions of wild-type and *Drp1*^−/−^ ESCs un-treated or treated with 1µM Thapsigargin for 6h. Graph shows cytochrome C protein levels normalized against ERK1/2 (cytosolic fraction) or ATP-b (membrane fraction). Average of 3 independent experiments +/-SEM is shown. Statistical analysis was done with a (C,E) Mann Whitney test *p<0.05, **p<0.01 or ***p<0.001.

**Supplementary Figure 4.**
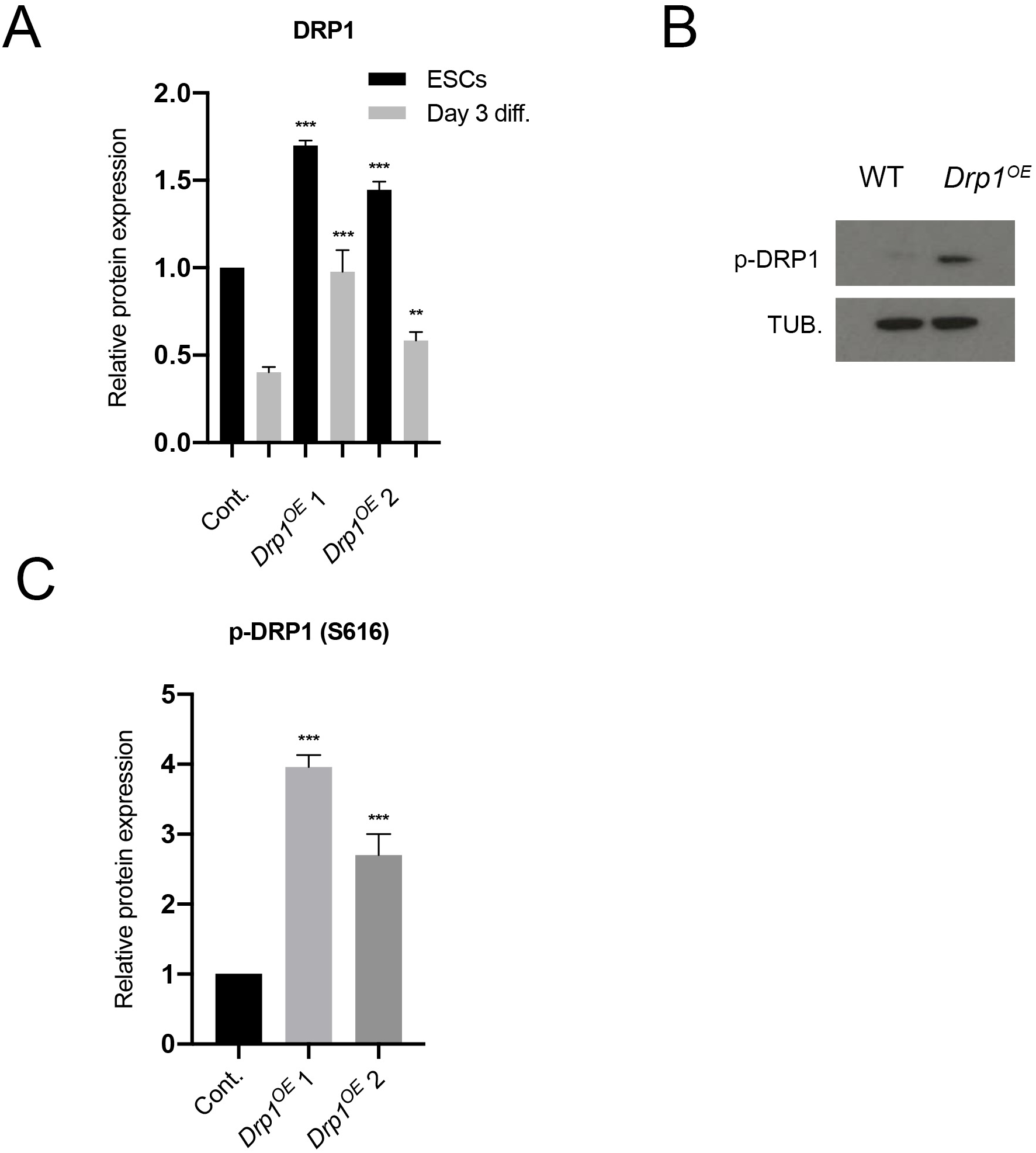
Characterization of *Drp1* over-expressing cells. **A.** Quantification of total DRP1 protein levels in wild-type and *Drp1*^*OE*^ ESCs and differentiating cells. Protein levels are normalized against α-TUBULIN (TUB.) and the graph shows expression levels relative to wild-type cells quantified from Figure 5A. **B.** Levels of phospho-DRP1 (S616) in wild-type and *Drp1*^*OE*^ cells at day 3 of differentiation in N2B27 detected by Western blot. **C.** Quantification of (B). Protein levels are normalized against αTUBULIN and the graph shows protein expression levels relative to wild-type cells. Average of 3 (C) or 5 (A) independent experiments +/-SEM is shown. Statistical comparisons are made to the control cells in the same culture condition. Students T-Test **p<0.01, ***p<0,001.

**Supplementary Figure 5.**
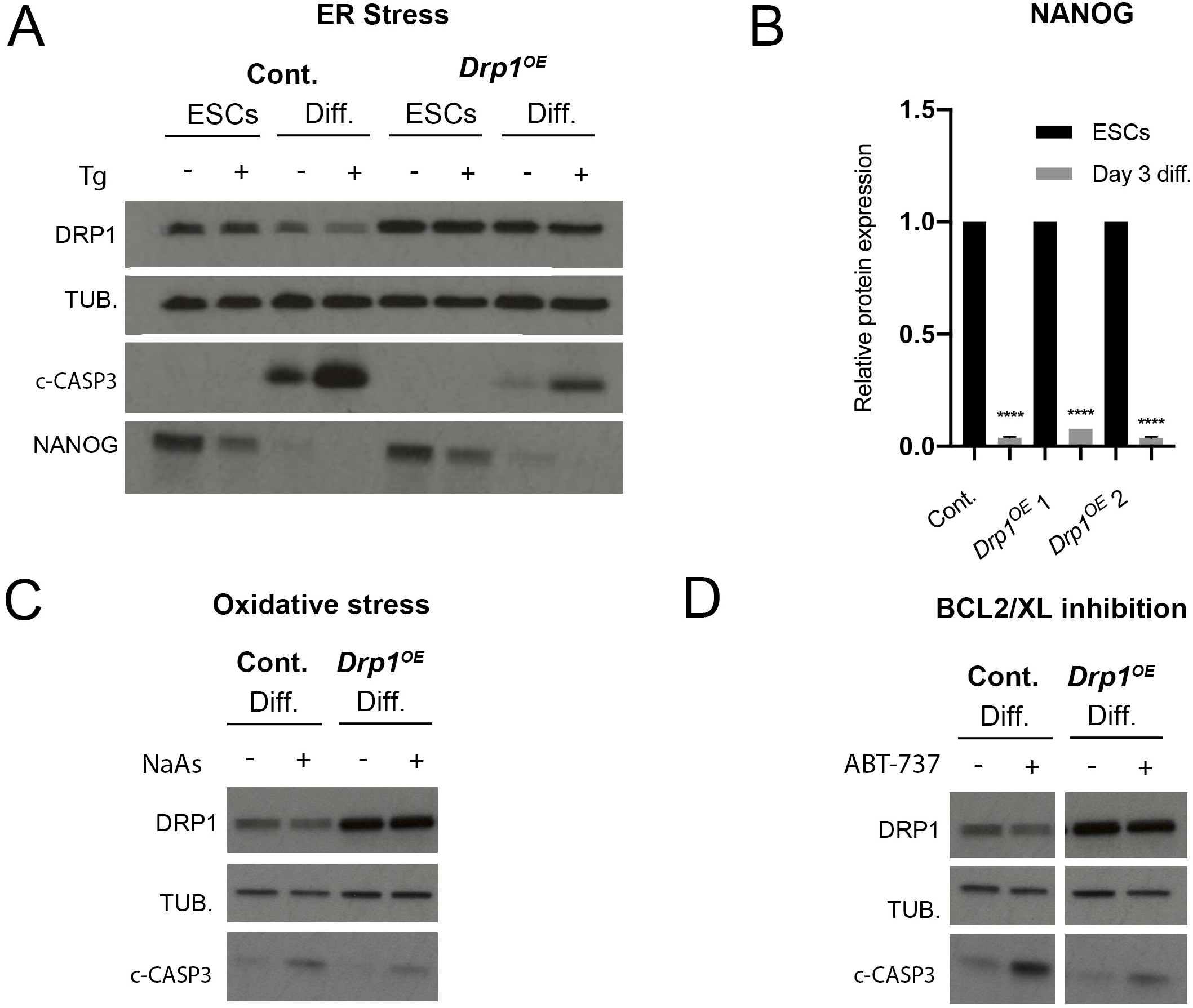
Stress response of *Drp1* over-expressing cells. **A.** Total DRP1, cleaved-CASPASE 3 (c-CASP3), α-TUBULIN (TUB.) and NANOG protein levels in wild-type and *Drp1*^*OE*^ ESCs at day 3 of differentiation in N2B27 untreated or treated with 1µM Thapsigargin for 5h. **B.** NANOG protein levels in wild-type and *Drp1*^*OE*^ ESCs and differentiated cells. Protein levels are normalized against α-TUBULIN and graph shows differentiation expression levels relative to ESCs for each cell type. **C.** Total DRP1, cleaved-caspase 3 and α-TUBULIN protein levels in wild-type and *Drp1*^*OE*^ at day 3 of differentiation in N2B27 untreated or treated with 1µM NaAs for 5h. **D.** Total DRP1, cleaved-caspase 3 and α-TUBULIN protein levels in wild-type and *Drp1*^*OE*^ at day 3 of differentiation in N2B27 untreated or treated with 1µM ABT-737 for 4h. The statistical comparisons are made to untreated cells. In B average of 3 independent experiments +/-SEM is shown. The statistical analysis compares a cell line in ESC and differentiation culture conditions. Students T-Test ****p<0.0001

## References

1. Bowling, S., Lawlor, K. & Rodriguez, T.A. Cell competition: the winners and losers of fitness selection. Development 146 (2019).

2. Snow, M.H.L. Gastrulation in the mouse: growth and regionalization of the epiblast. J. Embryol. exp. Morph. 42, 293–303 (1977).

3. Leitch, H.G., Tang, W.W. & Surani, M.A. Primordial germ-cell development and epigenetic reprogramming in mammals. Curr Top Dev Biol 104, 149–187 (2013).

4. Manova, K. et al. Apoptosis in mouse embryos: elevated levels in pregastrulae and in the distal anterior region of gastrulae of normal and mutant mice. Dev Dyn 213, 293–308 (1998).

5. Spruce, T. et al. An early developmental role for miRNAs in the maintenance of extraembryonic stem cells in the mouse embryo. Dev Cell 19, 207–219 (2010).

6. Heyer, B.S., MacAuley, A., Behrendtsen, O. & Werb, Z. Hypersensitivity to DNA damage leads to increased apoptosis during early mouse development. Genes Dev 14, 2072–2084 (2000).

7. Brown, E.J. & Baltimore, D. ATR disruption leads to chromosomal fragmentation and early embryonic lethality. Genes Dev 14, 397–402 (2000).

8. Dobles, M., Liberal, V., Scott, M.L., Benezra, R. & Sorger, P.K. Chromosome missegregation and apoptosis in mice lacking the mitotic checkpoint protein Mad2. Cell 101, 635–645 (2000).

9. Kalitsis, P., Earle, E., Fowler, K.J. & Choo, K.H. Bub3 gene disruption in mice reveals essential mitotic spindle checkpoint function during early embryogenesis. Genes Dev 14, 2277–2282 (2000).

10. Bolton, H. et al. Mouse model of chromosome mosaicism reveals lineage-specific depletion of aneuploid cells and normal developmental potential. Nat Commun 7, 11165 (2016).

11. Pernaute, B. et al. MicroRNAs control the apoptotic threshold in primed pluripotent stem cells through regulation of BIM. Genes Dev 28, 1873–1878 (2014).

12. Tait, S.W. & Green, D.R. Mitochondrial regulation of cell death. Cold Spring Harb Perspect Biol 5 (2013).

13. Prudent, J. & McBride, H.M. The mitochondria-endoplasmic reticulum contact sites: a signalling platform for cell death. Curr Opin Cell Biol 47, 52–63 (2017).

14. Chen, H. et al. Mitofusins Mfn1 and Mfn2 coordinately regulate mitochondrial fusion and are essential for embryonic development. J Cell Biol 160, 189–200 (2003).

15. Smirnova, E., Griparic, L., Shurland, D.L. & van der Bliek, A.M. Dynamin-related protein Drp1 is required for mitochondrial division in mammalian cells. Mol Biol Cell 12, 2245–2256 (2001).

16. Frezza, C. et al. OPA1 controls apoptotic cristae remodeling independently from mitochondrial fusion. Cell 126, 177–189 (2006).

17. Otera, H., Miyata, N., Kuge, O. & Mihara, K. Drp1-dependent mitochondrial fission via MiD49/51 is essential for apoptotic cristae remodeling. J Cell Biol 212, 531–544 (2016).

18. Estaquier, J. & Arnoult, D. Inhibiting Drp1-mediated mitochondrial fission selectively prevents the release of cytochrome c during apoptosis. Cell Death Differ 14, 1086–1094 (2007).

19. Frank, S. et al. The role of dynamin-related protein 1, a mediator of mitochondrial fission, in apoptosis. Dev Cell 1, 515–525 (2001).

20. Karbowski, M. et al. Spatial and temporal association of Bax with mitochondrial fission sites, Drp1, and Mfn2 during apoptosis. J Cell Biol 159, 931–938 (2002).

21. Breckenridge, D.G., Stojanovic, M., Marcellus, R.C. & Shore, G.C. Caspase cleavage product of BAP31 induces mitochondrial fission through endoplasmic reticulum calcium signals, enhancing cytochrome c release to the cytosol. J Cell Biol 160, 1115–1127 (2003).

22. Montessuit, S. et al. Membrane remodeling induced by the dynamin-related protein Drp1 stimulates Bax oligomerization. Cell 142, 889–901 (2010).

23. Renault, T.T. et al. Mitochondrial shape governs BAX-induced membrane permeabilization and apoptosis. Molecular cell 57, 69–82 (2015).

24. Lima, A., Burgstaller, J., Sanchez-Nieto, J.M. & Rodriguez, T.A. The Mitochondria and the Regulation of Cell Fitness During Early Mammalian Development. Curr Top Dev Biol 128, 339–363 (2018).

25. Stern, S., Biggers, J.D. & Anderson, E. Mitochondria and early development of the mouse. J Exp Zool 176, 179–191 (1971).

26. Nichols, J. & Smith, A. Naive and primed pluripotent states. Cell stem cell 4, 487–492 (2009).

27. Certo, M. et al. Mitochondria primed by death signals determine cellular addiction to antiapoptotic BCL-2 family members. Cancer Cell 9, 351–365 (2006).

28. Korsmeyer, S.J. Bcl-2 initiates a new category of oncogenes: regulators of cell death. Blood 80, 879–886 (1992).

29. Shepard, T.H., Muffley, L.A. & Smith, L.T. Mitochondrial ultrastructure in embryos after implantation. Hum Reprod 15 Suppl 2, 218–228 (2000).

30. Zhou, W. et al. HIF1alpha induced switch from bivalent to exclusively glycolytic metabolism during ESC-to-EpiSC/hESC transition. EMBO J 31, 2103–2116 (2012).

31. Knott, A.B., Perkins, G., Schwarzenbacher, R. & Bossy-Wetzel, E. Mitochondrial fragmentation in neurodegeneration. Nat Rev Neurosci 9, 505–518 (2008).

32. Sancho, M. et al. Competitive interactions eliminate unfit embryonic stem cells at the onset of differentiation. Dev Cell 26, 19–30 (2013).

33. Laurent, A. & Blasi, F. Differential DNA damage signalling and apoptotic threshold correlate with mouse epiblast-specific hypersensitivity to radiation. Development 142, 3675–3685 (2015).

34. Liu, J.C. et al. High mitochondrial priming sensitizes hESCs to DNA-damage-induced apoptosis. Cell stem cell 13, 483–491 (2013).

35. Csordas, G., Weaver, D. & Hajnoczky, G. Endoplasmic Reticulum-Mitochondrial Contactology: Structure and Signaling Functions. Trends Cell Biol 28, 523–540 (2018).

36. Booth, D.M., Enyedi, B., Geiszt, M., Varnai, P. & Hajnoczky, G. Redox Nanodomains Are Induced by and Control Calcium Signaling at the ER-Mitochondrial Interface. Molecular cell 63, 240–248 (2016).

37. Ishihara, N. et al. Mitochondrial fission factor Drp1 is essential for embryonic development and synapse formation in mice. Nat Cell Biol 11, 958–966 (2009).

38. Elgass, K., Pakay, J., Ryan, M.T. & Palmer, C.S. Recent advances into the understanding of mitochondrial fission. Biochim Biophys Acta 1833, 150–161 (2013).

39. Zhong, X. et al. Mitochondrial Dynamics Is Critical for the Full Pluripotency and Embryonic Developmental Potential of Pluripotent Stem Cells. Cell Metab 29, 979–992 e974 (2019).

40. Bahat, A. et al. MTCH2-mediated mitochondrial fusion drives exit from naive pluripotency in embryonic stem cells. Nat Commun 9, 5132 (2018).

41. Xie, Q. et al. Mitochondrial control by DRP1 in brain tumor initiating cells. Nat Neurosci 18, 501–510 (2015).

42. Kashatus, J.A. et al. Erk2 phosphorylation of Drp1 promotes mitochondrial fission and MAPK-driven tumor growth. Molecular cell 57, 537–551 (2015).

43. Nagdas, S. et al. Drp1 Promotes KRas-Driven Metabolic Changes to Drive Pancreatic Tumor Growth. Cell Rep 28, 1845–1859 e1845 (2019).

44. Brons, I.G. et al. Derivation of pluripotent epiblast stem cells from mammalian embryos. Nature 448, 191–195 (2007).

45. Guo, G. et al. Klf4 reverts developmentally programmed restriction of ground state pluripotency. Development 136, 1063–1069 (2009).

46. Nesterova, T.B. et al. Dicer regulates Xist promoter methylation in ES cells indirectly through transcriptional control of Dnmt3a. Epigenetics Chromatin 1, 2 (2008).

47. Schindelin, J. et al. Fiji: an open-source platform for biological-image analysis. Nature methods 9, 676–682 (2012).

48. Ramsby, M. & Makowski, G. Differential detergent fractionation of eukaryotic cells. Cold Spring Harb Protoc 2011, prot5592 (2011).

49. McCloy, R.A. et al. Partial inhibition of Cdk1 in G 2 phase overrides the SAC and decouples mitotic events. Cell Cycle 13, 1400–1412 (2014).

50. Ran, F.A. et al. Genome engineering using the CRISPR-Cas9 system. Nat Protoc 8, 2281–2308 (2013).

